# Single neuron diversity supports area functional specialization along the visual cortical pathways

**DOI:** 10.1101/2024.12.13.628359

**Authors:** M Feyerabend, S Pommer, MS Jimenez-Sosa, J Rachel, J Sunstrum, F Preuss, S Mestern, R Hinkel, M Mietsch, S Viyajraghavan, S Everling, S Treue, AFT Arnsten, DA Lewis, F Wolf, J Murray, Steve McCarroll, Lyle Muller, F Krienen, D Datta, XJ Wang, SJ Tripathy, G Gonzalez-Burgos, W Inoue, A Neef, JF Staiger, J Martinez-Trujillo

## Abstract

Humans and other primates have specialized visual pathways composed of interconnected cortical areas. The input area V1 contains neurons that encode basic visual features, whereas downstream in the lateral prefrontal cortex (LPFC) neurons acquire tuning for novel complex feature associations. It has been assumed that each cortical area is composed of repeatable neuronal subtypes, and variations in synaptic strength and connectivity patterns underlie functional specialization. Here we test the hypothesis that diversity in the intrinsic make-up of single neurons contributes to area specialization along the visual pathways. We measured morphological and electrophysiological properties of single neurons in areas V1 and LPFC of marmosets. Excitatory neurons in LPFC were larger, less excitable, and fired broader spikes than V1 neurons. Some inhibitory fast spiking interneurons in the LPFC had longer axons and fired spikes with longer latencies and a more depolarized action potential trough than in V1. Intrinsic bursting was found in subpopulations of both excitatory and inhibitory LPFC but not V1 neurons. The latter may favour temporal summation of spikes and therefore enhanced synaptic plasticity in LPFC relative to V1. Our results show that specialization within the primate visual system permeates the most basic processing level, the single neuron.

## Introduction

Human and other primates have hierarchically organized cortical visual processing streams composed of serially connected areas, starting in the primary visual area V1 and extending downstream to the lateral prefrontal cortex (LPFC, Felleman and Essen, 1991). The diversity of neuronal response patterns, selectivity to different visual stimuli, and involvement in diverse perceptual and cognitive functions across areas has been extensively documented in the macaque monkey (Merigan and Maunsell, 1993; Nassi and Callaway, 2009; Martinez-Trujillo, 2022). This inter-area functional diversity has been implicitly assumed to be due to variations in expression levels of receptors subtypes, synaptic strength, inputs, and connectivity within an area’s microcircuit (Carandini and Heeger, 2011; Godlove et al., 2014; Ardid et al., 2015; Jiang et al., 2015). It has been further proposed that neurons across cortical areas of mammals follow the principle of serial homology, i.e., single neuron classes are small quantitative variations of a common theme (Harris and Shepherd, 2015). Here we test the alternative hypothesis that single neurons in different visual areas of primates show qualitative and quantitative variations in their intrinsic morphological and functional properties that contribute to inter-area specialization.

Previous studies have shown that neurons in macaque area V1 show little changes in their tuning for basic visual features during training, even when subjects’ show a strong improvement of behavioral discrimination thresholds for the same features (Ghose et al., 2002). On the other hand, neurons in the lateral intraparietal area (LIP) downstream from V1 acquire selectivity for categories during training (Freedman and Assad, 2006). In the LPFC, downstream from LIP, neurons flexibly become selective for new associations of visual features during intervals of minutes or hours (Rouzitalab et al., 2023; Abbass et al., 2024). At the molecular level, a recent study in macaques reported that the ratio of NMDA to AMPA receptors in synapses is larger in the LPFC than in area V1 suggesting that LPFC circuits support learning via synaptic plasticity to a larger degree than V1 circuits (Yang et al., 2018). Thus, the primate visual pathways extend along a cortical *stability – plasticity* axis: Neurons in early areas are *stably* tuned to basic visual features and neurons in far downstream areas *flexibly* ‘learn’ novel feature combinations encountered in new situations or environments. One issue that remains poorly investigated is whether variations in cells’ intrinsic properties and morphology could shape single neuron computations and synaptic plasticity, ultimately contributing to inter-area specialization along this axis.

Reports of intrinsic electrophysiological properties and morphology of neurons in primates are scarce relative to other species such as the mouse. Previous studies showed that excitatory pyramidal neurons in area V1 and the LPFC of macaques diverge in their morphology (Amatrudo et al., 2012; Medalla and Luebke, 2015; Gilman et al., 2017), whereas pyramidal cells in mouse areas V1 and dorsal PFC do not have such prominent differences (Gilman et al., 2017). Moreover, a study in macaques has revealed differences in the response patterns of pyramidal cells between the lateral intraparietal (LIP) area and the LPFC (González-Burgos et al., 2019). It is possible that not only excitatory but also inhibitory neurons have diversified their intrinsic properties to contribute to the functional specialization of cortical areas in the primate visual pathways. However, we have not found any study systematically comparing intrinsic properties of inhibitory interneurons between different areas of the primate visual processing hierarchy.

We investigate this issue in the common marmoset (*Callithrix jacchus*), a non-human primate (NHP) model that has become popular amongst neuroscientists due to their faster reproductive cycle, amenability to transgenic manipulations and similarity of visual processing with humans (Mitchell and Leopold, 2015; Okano et al., 2016). We compare intrinsic electrophysiological properties and morphology of single neurons between areas V1 and the LPFC of marmosets. We use methods, protocols and tools similar to those the Allen Institute of Brain Science (AIBS) used in the mouse and human cell type database (Gouwens et al., 2019), i.e. whole-cell patch clamp recordings in acute slices and morphological reconstructions of single neurons. We built a “Primate Cell Type Database” resource (https://primatedatabase.com). It contains 374 biophysical characterizations of intrinsic membrane properties as well as morphological reconstructions of neurons from several areas (predominantly V1 and LPFC). We found that intrinsic properties and morphology of excitatory and inhibitory neurons quantitatively and qualitatively varied between V1 and the LPFC. Thus, area functional specialization along the primate visual cortical pathways occurs at the most basic level of processing, the single neuron.

## Materials & Methods

### Statement on animal research

Research on NHPs is an ethically sensitive but irreplaceable part of neuroscience research. The authors of this study are committed to pursue the best scientific result with the least possible harm to the animals. Data was collected from 42 (male: 31, female: 11) common marmosets (*Callithrix jacchus*, 26 at Western University + 16 at University of Göttingen) with a median age of 4.43 years (IQR: 4.81, min: 1.52, max 18.06). Animal research from the Canadian group was conducted in accordance with the Canadian Council of Animal Care policy on the care and use of laboratory animals. The experiments were approved by the Animal Care Committee of the University of Western Ontario. Animals from the German research group were fostered and kept at the German Primate Center, Göttingen, Germany. Husbandry and experiments were conducted in compliance with the Directive 2021/63/EU of the European Union and the German Animal Welfare Act and therefore meet the regulations of the European Animal Research Association. The animals were sacrificed as part of a broad study at the German Primate Center approved by the Lower Saxony State Office for Consumer Protection and Food Safety (LAVES; reference number 33.19-42502-04-20/3458). Experimental procedures were in accordance with regulations of the German Animal Welfare Act. All animals were under constant veterinary supervision.

### Acute marmoset ex vivo brain slice preparation

Marmosets were anaesthetized with Ketamine (20 mg/kg, intramuscular) and isoflurane (2–5 %) and then euthanized by trans-cardial perfusion (Canada) or pentobarbital (Germany). In the latter case, ice-cold solution was poured over the head before opening the skull to slow down cellular processes and decrease brain deterioration during organ removal. Afterwards, the brain was immediately rinsed with pre-chilled (2–4 °C) N-methyl-D-glucamine (NMDG) substituted artificial cerebrospinal fluid (aCSF) containing 92 mM NMDG, 2.5 mM KCl, 1.2 mM NaH_2_PO_4_·H_2_O, 30 mM NaHCO_3_, 20 mM 4-(2-hydroxyetyl)-1-piperazineethansulfonic acid (HEPES), 25 mM glucose, 5 mM Na-ascorbate, 2 mM thiourea, 3 mM Na-pyruvate, 10 mM MgSO_4_·7H_2_O and 0.5 mM CaCl_2_·2H_2_O. The pH was adjusted to 7.3–7.4 with concentrated hydrochloric acid (37 %) and the osmolality was set to 300–305 mOsm/kg. The solution was oxygenized with 95 % O_2_ and 5 % CO_2_ (carbogen) for 15 minutes prior to animal surgery. The brain was transferred into a container containing the same NMDG aCSF and quickly transported from the animal facility to the laboratory room (10–20 min) (Ting et al., 2018; Gouwens et al., 2019).

The hemispheres were separated and trimmed to blocks of prefrontal and visual cortex. Slices of 300 µm were cut with a vibratome VT1200 S (Leica) using the same ice-cold NMDG aCSF as above. Subsequently, the slices were transferred to a recovery chamber filled with NMDG aCSF at 32 °C and incubated for 12 minutes before stored in a holding chamber at room temperature. This chamber was filled with HEPES aCSF containing 92 mM NaCl, 2.5 mM KCl, 1.2 mM NaH_2_PO_4_·H_2_O, 30 mM NaHCO_3_, 20 mM HEPES, 25 mM glucose, 5 mM Na-ascorbate, 2 mM thiourea, 3 mM Na-pyruvate, 2 mM MgSO_4_·7H_2_O and 2 mM CaCl_2_·2H_2_O. The slices were stored for up to 10 hours with minimal submersion and transferred to fresh HEPES aCSF after 6 hours. HEPES provided additional support to pH buffering and reduced slice deterioration over extended time periods. All solutions were continuous oxygenized with carbogen gas and set to 300–310 mOsm/kg.

All solutions for ex vivo brain slice preparation and electrophysiological recordings were adopted from the technical white paper from the AIBS Cell Type Data base. The latter facilitates the comparison between our marmoset dataset with mouse and human datasets from AIBS Cell Type Data base (Gouwens et al., 2019; Berg et al., 2021).

### Patch clamp electrophysiology

Individual brain slices were transferred into the recording chamber and continuously perfused with ∼32 °C carbogen-saturated aCSF containing 126 mM NaCl, 2.5 mM KCl, 1.25 mM NaH_2_PO_4_·H_2_O, 26 mM NaHCO_3_, 20 mM HEPES, 12.5 mM glucose, 1 mM MgSO_4_·7H_2_O, 2 mM CaCl_2_·2H_2_O and a mix of synaptic blockers (2 mM kynurenic acid and picrotoxin 0.1 mM). Thick-walled borosilicate glass pipettes were manufactured on the day of recording and filled with intracellular solution containing 126 mM K-gluconate, 4 mM KCl, 0.3 mM EGTA, 10 mM HEPES, 4 mM K_2_-ATP, 0.3 mM Na-GTP, 10 mM Na_2_-Phosphocreatinine. The pH was adjusted to 7.3–7.4 with 1 M KOH and the osmolality was set to 290 mOsm/kg with sucrose when necessary. The internal solution was supplemented with 0.5% biocytin to fill the recorded neurons for histology. Pipette resistance in the bath was 3-7 MΩ. Recording equipment varied across experimenters: data were acquired with either a Multiclamp 700B and a Digidata (Axon instruments/ molecular devices), a SEC-10LX or SEC-05X (npi electronics) digitized with a 1401-3A (CED) or an EPC 10 USB (HEKA). After formation of a stable seal above 1 GΩ and break-in, access resistance was determined in voltage clamp (VC) mode. Then, the recording mode was switched to current clamp (CC) and bridge balance and pipette neutralization were applied. When necessary, a holding current was applied and continuously readjusted throughout to maintain a membrane potential close to the initial value after break-in. Cells that required more than 20 pA holding current to keep the membrane potential at -60 mV in the initial whole-cell configuration were generally not recorded from. Recording protocols and procedures were standardized across laboratories and acquisition equipment. Neurons were subjected to hyper- and depolarizing 1 s square pulse current injections. Furthermore, 275 (73.5%) cells were recorded with an additional 3 ms short square pulse protocol.

### Histology

After recording, slices were fixed with 4% m/v PFA + 15% v/v picric acid (1.3% saturated in water) at 4 °C for a period of 24–36 hours and then washed in PBS. Then, slices were permeabilized by incubation in PBS + 2% Triton-X for 2 x 15 min. Biocytin filling of patched cells was made visible by a 4-hour incubation of streptavidin conjugated with Alexa Flour 647. After a subsequent washing step, slices were incubated in PBS with DAPI (1:4000), before mounted on specimen slides with aqua-polymount.

Imaging was done using a SP8 confocal microscope (Leica, Germany) or a LSM 880 with Airyscan (Zeiss, Germany) to assess the cell location, quality of the filling and dendritic type. Dendritic type was determined based on spininess and somatodendritic configuration of the cell similar to Gouwens et al. 2018. Cells with sufficiently retained dendritic and axonal arborization were used for another acquisition with 40x or 63x magnification (z-step size 0.5–0.8 µm). These image stacks were then used to reconstruct the neuronal morphology using Neurolucida (MBF Bioscience). Automated quantifications of neurites were obtained with Neurolucida explorer (MBF Bioscience, USA).

### Analysis of electrophysiological features

Raw recordings were converted into the neurodata without borders (nwb) format (version 2.4.0; (Rübel et al., 2022)) using the MATNWB repository (https://neurodatawithoutborders.github.io/matnwb/). Additional information such as subject data (age, sex, etc.), acquisition parameters (access resistance, temperature, etc.) or histological specifications (layer, putative superclass, etc.) were added if available. A custom MATLAB analysis pipeline was used for quality control and extraction of electrophysiological features. The code is publicly available on GitHub (<u>GitHub - mfeyerab/MATFX</u>). Recordings from cells were not considered for further analysis if one of the following criteria was met: (1) an initial membrane potential above -55 mV or (2) input resistance or action potential (AP) waveform features could not be determined. In addition, recording quality was assessed in a sweep-wise fashion. For this purpose, a time window of 250 ms for each pre and post stimulus period was selected. A sweep was discarded if: (1) the membrane potential in either segment exceeded a root mean square error of 0.65, (2) the membrane potential difference between pre and post stimulus exceeds 2.5 mV, (3) the membrane potential in the pre-stimulus segment is not more depolarized than -52.5 mV, (4) the membrane potential of the pre-stimulus deviates more than 6 mV from the median pre-stimulus membrane potential of all passing sweeps, (5) flagged by manual curation based on evaluation of the pre-stimulus test pulse (see **Table 1** and **STab 1 for a list of key features, and STab 2** for description of all electrophysiological features).

**Table 1:** Electrophysiological characteristics of class 3 neurons in LPFC and V1. Comparison between V1 and LPFC in key electrophysiological features of C3 Cells. The measure of central tendency of the distribution median. Negative efect size measures show smaller values for V1 compared to PFC. p Values are corrected with Benjamini-Hochberg procedure.

| Feature | LPFC<br>n = 104 | V1<br>n = 56 | adj. p-value | z-val. | Effect size [CI] |
| --- | --- | --- | --- | --- | --- |
| Vrest, mV | -71.61 | -71.68 | 0.6721 | 0.47 | -0.04 [-0.19; 0.11] |
| R<sub>inHD</sub>, MΩ | 212.6 | 461.4 | <1e-4* | -7.80 | 0.62 [0.53; 0.70] |
| rheobase, pA | 90 | 30 | <1e-4* | 7.95 | -0.63 [-0.71; -0.54] |
| tau, ms | 26.74 | 30.28 | 0.2640 | -1.18 | 0.09 [-0.06; 0.25] |
| sag ratio | 1.13 | 1.21 | 0.0012* | -3.45 | 0.27 [0.11; 0.43] |
| inst. rectification | 0.92 | 0.75 | <1e-4* | 8.08 | -0.64 [-0.73; -0.54] |
| AP width | 0.35 | 0.27 | <1e-4* | 5.28 | -0.42 [-0.54; -0.29] |
| amp slow trough | -19.65 | -21.65 | 0.0127* | 2.60 | -0.21 [-0.36; -0.05] |
| rate of rheo, Hz | 1.00 | 1.00 | 0.1274 | -1.64 | 0.13 [-0.03; 0.30] |
| med. inst. rate, Hz | 22.2 | 19.3 | 0.0116* | -2.65 | 0.22 [0.07; 0.37] |
| IQR inst. rate, Hz | 7.30 | 8.10 | 0.2234 | 1.32 | -0.11 [-0.27; 0.05] |
| adaRat Blast/B1 | 0.225 | 0.048 | 0.0032* | -3.14 | 0.25 [0.11; 0.39] |
| flslope, Hz/pA | 0.048 | 0.123 | <1e-4* | -5.07 | 0.41 [0.26; 0.53] |
| cvISI | 0.354 | 0.249 | 2.25e-4* | 3.90 | -0.34 [-0.49; -0.17] |
| rate of hero, Hz | 8 | 13 | 0.0116* | -2.68 | 0.21 [0.06; 0.36] |
| latency at hero, ms | 14.35 | 16.41 | 0.9544 | 0.06 | 0.00 [-0.17; 0.16] |
| adaptation index | 0.091 | 0.048 | <1e-4* | 4.51 | -0.40 [-0.55; -0.25] |
| difference in trough | 4.608 | 6.760 | 0.0053* | -2.96 | 0.24 [0.09; 0.39] |
| burst index | 0.888 | 0.782 | <1e-4* | 4.48 | -0.39 [-0.52; -0.24] |

**Table 2:**
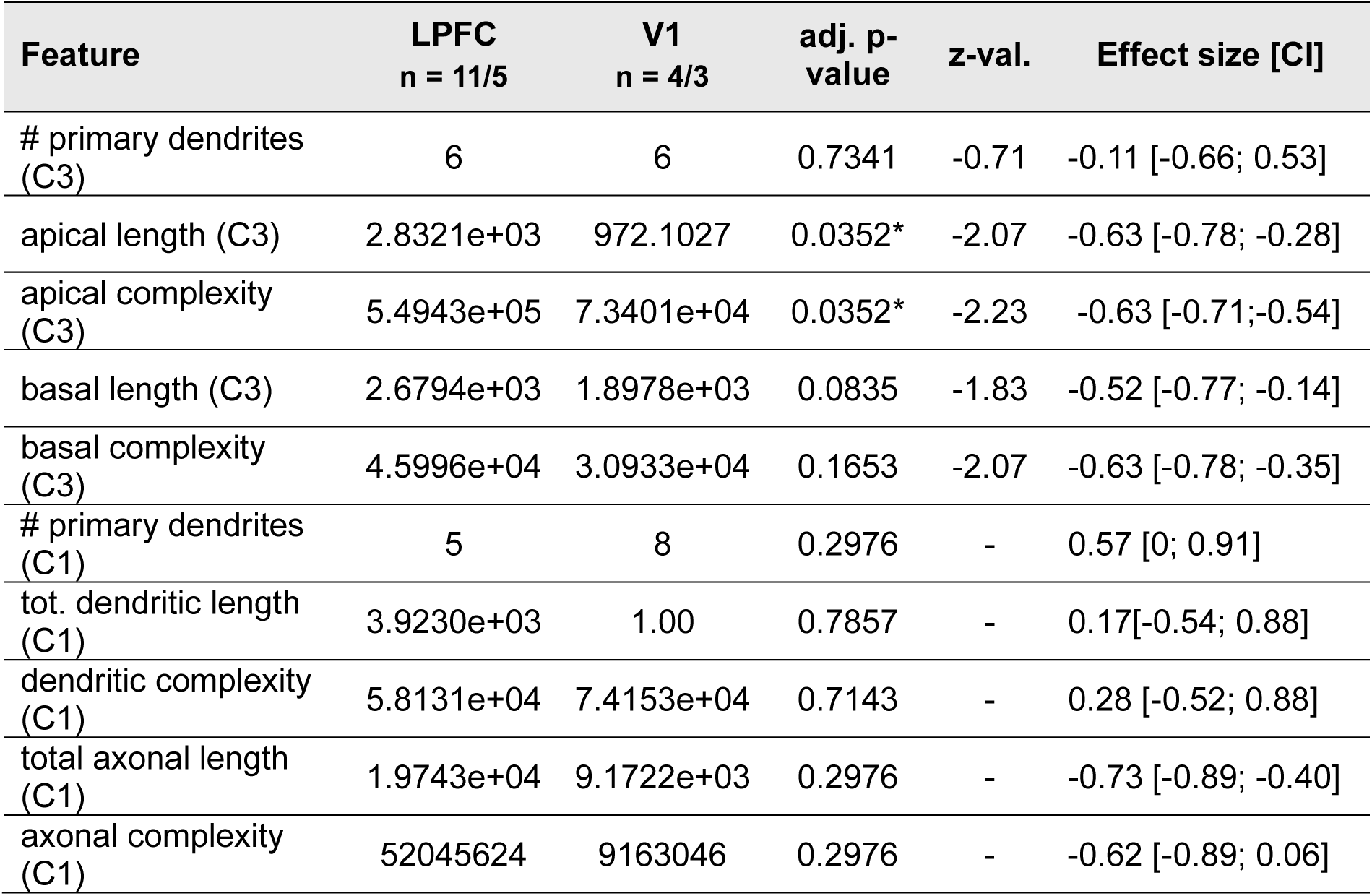
Comparison between V1 and LPFC in key morphological features of class 1 and 3 neurons. Values are median of electrophysiological characteristics in class 1 cells of LPFC and V1. Efect sizes were calculated as rank-biserial correlation. Negative values indicate larger LPFC values.

### UMAP and cross-species classification

Uniform Manifold Approximation Projection, UMAP (McInnes et al., 2018) space was created by combining three data sets each obtained from a different species. Mouse and human data were available at the AIBS Cell Type Database. For the classification analyses separate training and test data sets for each species were chosen. Cells with a missing value in more than 15% of features were excluded from the subsequent analysis yielding a total number of 2421 cells (1757 mouse, 356 human, and 308 marmoset). Initially, 29 electrophysiological features (see supplement for more detail) were reduced to 16 components by probabilistic PCA explaining more than 95% of the total variance. Then data was transformed into UMAP space (McInnes et al., 2018) using custom MATLAB scripts in two instances: (1) in a space with two components for visualization of electrophysiological diversity in the different data sets (number of nearest neighbours = 60, minimum distance = 0.2) and (2) in a space with three components for supervised classification across species (number of nearest neighbours = 30, minimum distance = 0.3).

For classification, three labels were chosen: Class 1 (C1), which contains fast-spiking interneurons (FSI); Class 2 (C2), which contains the non-fast spiking population of interneurons and Class 3 (C3), which contains excitatory cells (EXC). Ground truth data (**SFig. 1B**) was obtained by Cre reporter lines and electrophysiological types (e-types, determined by unsupervised clustering by Gouwens et al. 2019), to exclude contamination by off-target expression (Hu et al., 2013). Cells which expressed Cre with the parvalbumin (PV) promoter and an e-type ranging from Inh_8 to Inh_13 were considered fast spiking interneurons (FSI). Cells which had a spiny dendritic type were considered EXC. Cells that expressed Cre with a somatostatin (SST) promoter and an e-type ranging from Inh_4 to Inh_7 were considered SST cells, whereas cells that expressed Cre with a serotonin receptor 3a (5-HT3A) or vasoactive (poly)peptide (VIP) promoter and had either e-type 2 or 3 were considered VIP cells. Lastly, cells that had the e-type “Inh_1”, which was marked by low spike-frequency adaptation and commonly high latency of the rheobase spikes and expressed Cre with either a 5-Htr3a or a neuron-derived-neurotrophic-factor (Ndnf) promoter were considered neurogliaform (NGF) cells. C2 consisted of the SST, VIP and NGF subtype, which do not show the distinctive fast spiking phenotype of PV cells. Cell type composition of training data was informed by histological data of the primate neocortex, consequently the ratio of SST, VIP and NGF cells within the training data was fixed to 2:2:1. A random forest classifier for cross species cell labeling was trained (Seiffert et al., 2010) with uniform priors with a subset of the 3 component UMAP data consisting of 1174 mouse cells with marker labels. Cell composition of test data was matched across species and it was biased towards excitatory cells (∼65 %). Marmoset and human test data set for cross-species validation was determined by unambiguous morpho-electrophysiological identity (see examples in **Fig 2** and **SFig 1**).

### Statistical analysis

All hypothesis testing regarding differences between areas was done with 2-sided rank sum tests in MATLAB and effect sizes were reported by rank bi-serial correlation (r_rb_). All r_rb_ values and their 95% confidence intervals were calculated with the measures-of-effect-size-toolbox (Hentschke and Stüttgen, 2011). Reported p-values were corrected for false discoveries rates via the Benjamini-Hochberg procedure. Correlations reported were either tested by Spearman (correlation coefficient: ρ) or Pearson (correlation coefficient: r).

The residual values of rheobase were obtained from a linear regression model with rheobase as dependent variable and the ratio between the difference in the action potential threshold and the resting membrane potential (Dvoltage), and the input resistance (R) as predictor variable. This model (rheobase = B0 + B1*(Dvoltage/R) + residuals) captures the passive/ohmic aspect of the rheobase.

### Effect of different acquisition systems on AP waveform

Our data were recorded at two different sites with different acquisition systems. Therefore, we checked whether this had an effect on electrophysiological measurements. Data recorded with the HEKA amplifier show significantly higher AP amplitudes than those recorded with npi (**SFig 2A**), likely because differences in pipette capacitance compensation between amplifiers. Uncompensated pipette capacitance, together with the access resistance, create a passive low-pass filter which artificially shorten and widen the AP. This effect was stronger in neurons with narrow spike waveforms. Selection of AP waveform features for the UMAP procedure was informed by these observations. AP width was the only wave form parameter chosen as input, since it showed no significant effect between the amplifier systems (**SFig 2B**). Ultimately, neurons in the UMAP projection did not suggest clustering by acquisition equipment (**SFig 2C**) and we conclude that our intrinsic property profiles are mainly shaped by biological factors such as cell identity.

## Results

We recorded from a total of 463 neurons in slices obtained from 42 marmosets. 374 neurons (80.8%) passed our quality control (QC pipeline) to enter the primate cell type database (see **Fig 1**). This database contains 107 V1 cells, 256 LPFC cells (focused on Brodmann area (BA) 8 and medial part of BA 46) and 11 cells from cingulate cortex (not included in this study). We targeted both pyramidal and non-pyramidal cells focusing on supragranular layers. A total of 144 neurons were examined histologically, and spine density was evaluated to classify the cells into aspiny, sparsely spiny or spiny. The latter could further inform whether a neuron was inhibitory or excitatory (Gouwens et al., 2019). 29% of the cells evaluated were either aspiny or sparsely spiny (for clarity purposes we will refer to the latter two categories as aspiny), and 71% were considered spiny. The marmoset dataset contains a wide spectrum of morphologically different cell types (**Fig. 2A, B**). Of the 144 neurons evaluated morphologically, a total of 32 cells were reconstructed. We also used two other data sets from mouse and human, respectively, which are publicly available from the Allen Institute for Brain Science’s (AIBS) Cell Type Database (Gouwens et al., 2019). After applying the same QC pipeline as in the marmoset data, we included 1907 of 1920 AIBS mouse cells (99.32%) and 403 of 413 human cells (97.58%) in our analyses **(SFig 1A)**.

**Figure 1:**
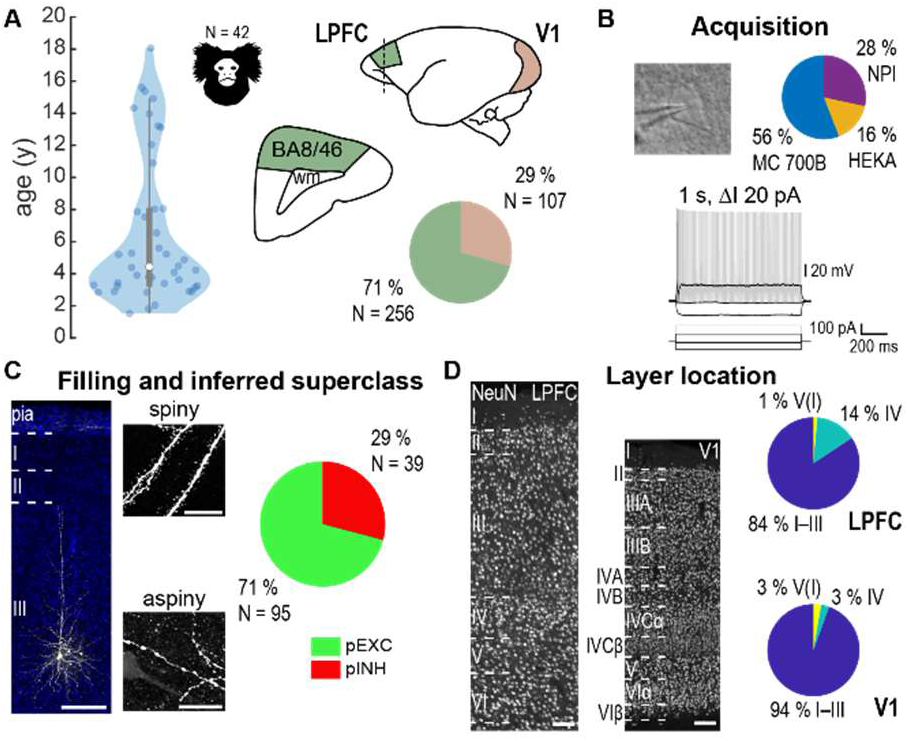
Tissue acquisition and characterization of neuronal morphology and electrophysiology in multiple areas of the marmoset neocortex. **A:** Acute slices of LPFC and V1 were obtained from 42 marmosets with a median age of 4.4 years. Neurons were characterized by electrophysiology (B) and morphology (C). **B:** Cells (grayscale picture) were subjected to 1 sec long square pulse current injection with a 20 pA increment from -110 pA till suprathreshold saturation. Example traces show hyperpolarizing, subthreshold, rheobase and suprathreshold injections. Data was acquired across different laboratories and recording systems (right pie chart). **C, D:** Anatomical evaluation of all filled neurons consisted of determination of broad class (spiny, putative Excitatory, pEXC versus aspiny, putative Inhibitory, pINH) via somatodendritic configuration, spininess (C) and cortical home layer of the soma (D). Left image in C shows a biocytin filled pyramidal neuron in layer III of LPFC; nuclei are stained with DAPI (blue); scale bar equals 150 µm. Small images show example spiny (top) and aspiny (bottom) somatodendritic morphology; scale bars equal 20 µm. Pie-chart shows the distribution of morphologically identified cells. In D the image shows NeuN staining with layer delineation in LPFC and V1; scale bars equal 100 µm. Right pie-charts represent the distribution of cells by layers for LPFC and V1. Cells recorded in both areas were predominantly located in supragranular (I-III) layers.

**Figure 2:**
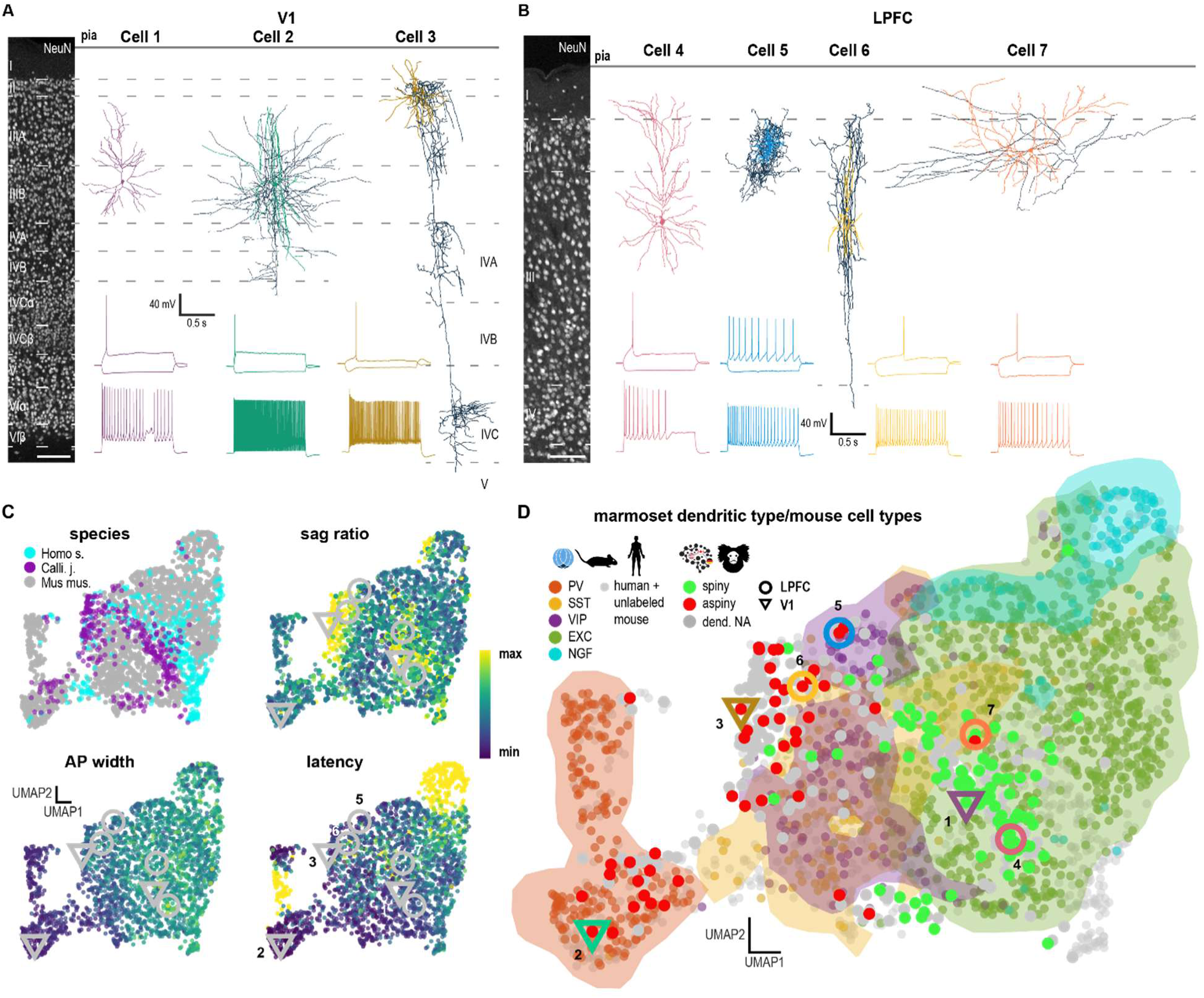
Neuronal diversity within marmoset data and co-projection with known cell types. **A, B:** Examples of different marmoset cells separated by cortical area (A: V1, B: LPFC). Cells 1 and 4 are regular-spiking pyramidal cells. Cell 2 is a fast-spiking basket cell. Cell 3 is a vertically oriented basket cell. The cell was recorded in the dorsal medial V1 with expanded layers IV–VI. Cell 5 is a neurogliaform cell. Cell 6 is a double bouquet cell and cell 7 is a regular spiking spiny non-pyramidal cell (Below each cell is the corresponding subthreshold, rheobase and hero sweep. Scale bars equal 100 µm. **C:** Panel of 2D UMAP projections visualizing distribution of cells by species and key electrophysiological features. **D:** 2D UMAP projection visualizing distribution of marmoset cells color-coded by dendritic type, referenced with known cell type in mouse. Colored background delineates areas with high density for the respective cell type. Location of examples from subfigures A and B are indicated by their respective color (downward facing triangle: V1, circle: PFC). The Logos indicate the origin of the data. Allen Institute in blue/black. Our NEURONEX consortium in black + country flags.

### Comparing profiles of intrinsic properties across species

We combined our marmoset data with the two AIBS datasets in a multivariate map of electrophysiological features to determine whether the cell type profiles of intrinsic membrane properties of marmoset, mouse and human cortex neurons overlap under varying factors such as species or cortical area (**Fig 2C**). Importantly, our data were acquired with protocols and solutions identical to those used in the AIBS studies, thus minimizing the influence of methodological factors on potential inter-species differences (see Methods). Intrinsic properties of all cells were quantified in the same analysis pipeline by 29 electrophysiological features covering the subthreshold domain, action potential (AP) shape, and AP (spike) train pattern diversity (see **STab 1** for a list and respective quantification of features). Subsequently, we used the dimensionality reduction procedure “Uniform Manifold Approximation and Projection” (UMAP) for visualizing intrinsic properties profiles (**Fig 2C**). While the data showed a general overlap across species, marmoset and human cells were concentrated in hot zones marked by large sag: a delayed deflection in the membrane potential response to hyperpolarizing current injections (**Fig. 2C** upper right panel, mouse data indicated by gray filled circles in **Fig. 2C** upper left panel).

To better understand the distribution of different cell types on the UMAP projection, we utilized available cell type labels from 1460 mouse cells (76.6 % of analyzed mouse cells). Neurons showing the “spiny” dendritic type were considered to be excitatory and 4 major interneuron subtypes (PV/SST/VIP/NGF) were identified by a combination of e-type (Gouwens et al., 2019) and labeling by one of five Cre-lines (PV-Cre/SST-Cre/VIP-Cre/5HT3A-Cre/NDNF-Cre; for more detail see **SFig 1**). High density regions of the corresponding cell identity are indicated by color shading in **Fig 2D** (see also **SFig 1**). One of the most prominent differences between GABAergic and EXC cells is the speed of repolarization after the peak of an AP, with inhibitory cells having a narrower AP. Indeed, the first UMAP component is strongly influenced by AP width (r = 0.745, p < 0.001, see gradient in **Fig. 2C** lower panel) with FSI cells clustering to the left of the map. This electrophysiological profile is strongly associated with PV fast spiking interneurons, which was in line with the corresponding cell type cluster (see orange background colors of **Fig. 2D**). Although different cell types tend to localize to a certain region in the UMAP projection, they rarely separated into well-defined clusters but rather occupied a certain segment on a continuous spectrum of intrinsic properties. VIP and SST cells, indicated by purple and gold, showed a high degree of overlap.

The dendritic type of marmoset neurons (spiny vs. aspiny) showed a good correspondence with mouse cell types, with occasional intermixing between aspiny and spiny cells (see **Fig 2D** and **SFig 3B** for dendritic type of marmoset cells). The only notable shift of intrinsic property profiles could be observed for neurogliaform (NGF) cells, which in mouse were associated with low spike frequency adaptation and commonly high spike latencies (**Fig. 2D** cyan marker and background). Interestingly, all 6 morphologically identified NGF cells in marmoset showed strong spike frequency adaptation and short spike latencies (see **SFig 4**). Except for NGF cells, these results suggest that overall feature profiles of distinct cell types overlap across the different species.

### Classification of marmoset cell types based on intrinsic properties

We merged the four interneuron subpopulations of AIBS mouse data (i.e., PV, SST, VIP, NGF) into two group labels, labelling the PV cells were labeled as class 1 (C1), while the SST, VIP, and NGF were grouped as class 2 (C2). The label EXC was preserved, making class 3 (C3). We projected the intrinsic electrophysiological properties onto a 3D UMAP (**Fig 3**). We trained a random forest classifier (Seiffert et al., 2010) to predict the class label (C1, C2, C3) from the UMAP components using a sample of 1174 mouse neurons. The procedure was repeated 500 times with random training sets of fixed numbers of cells (**Fig. 3A**). We finally used a sample of 150 cells (not included in the training set) to assess the classifier performance in the mouse data. Median test performance in the mouse across the different classes was 91.3% (**Fig. 3B**).

**Figure 3:**
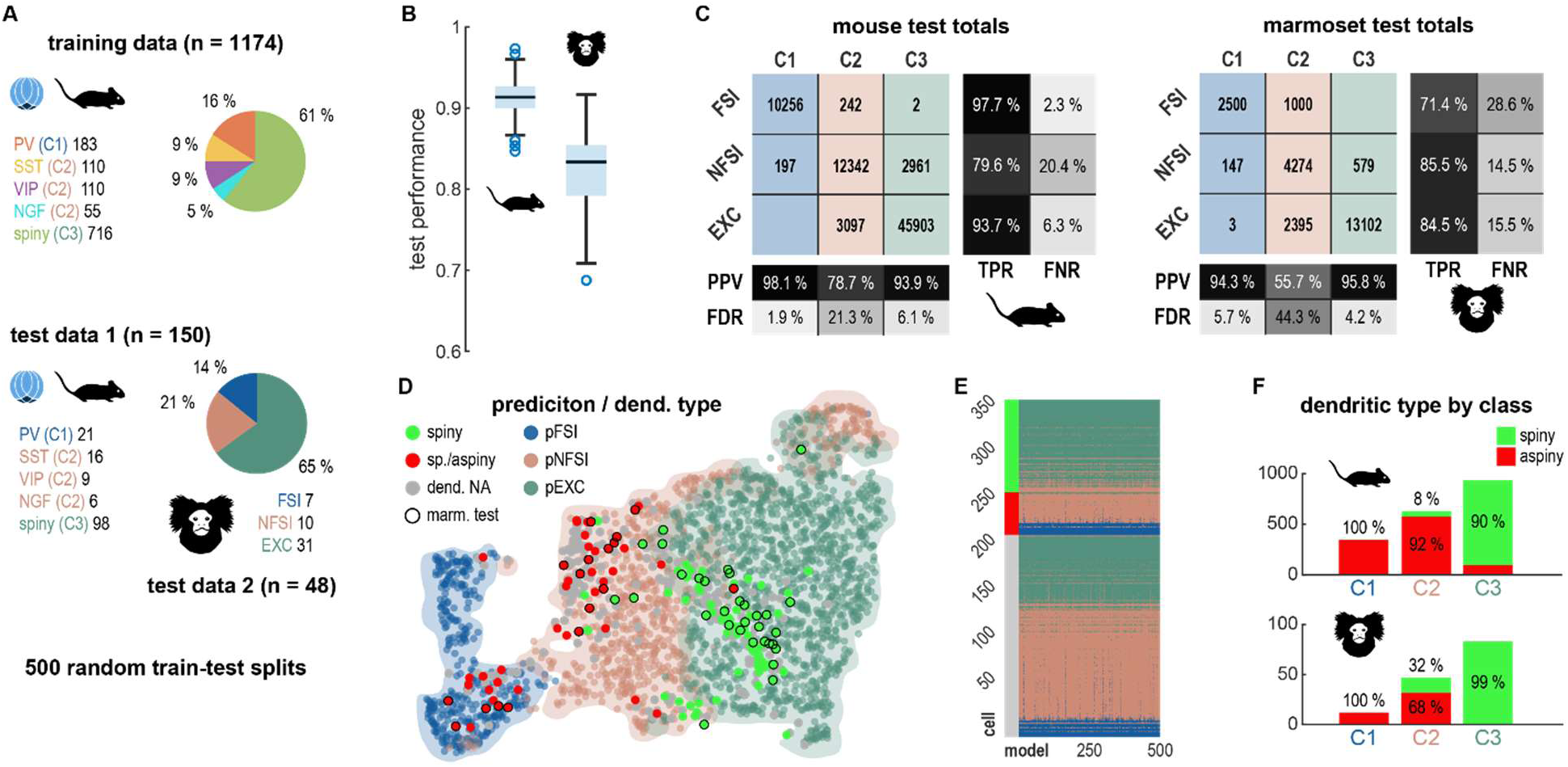
Objective classification of primate cells. **A:** Panel of pie charts showing cell type composition of training and test data. Three different classes were used: Class 1 (dark blue; C1), Class 2 (ocher, C2), merged from SST (yellow), VIP (purple) and NGF (turquoise) cells and Class 3 (dark green, C3). Mouse and marmoset test data had identical proportion of classes. For marmoset test data we chose electrophysiology with congruent morphology for C1 and C2, respectively. Marmoset C3 cells were drawn from spiny cells. The procedure was repeated 500 times with randomly drawn training and test sets. **B:** Performance of classifier across repetitions. Horizontal boxplots show distribution of performances. Median test performances by species were 91.3% for mouse and 83.3 % for marmoset. C: Confusion matrices for mouse and marmoset test data showing all classification totals. The positive predictive value (PPV) for C1 and C3 cells is over 90% in the marmoset. FDR = False Discovery Rate, TPR = True Positive Rate, FNR = False Negative Rate. **D:** Example UMAP visualization of one model prediction referenced with dendritic type of marmoset cells. Shaded areas indicate high density of the respective class. **E:** Colormap visualizing prediction outcome of marmoset cells for all 500 repetitions. Cells are grouped by dendritic type and then sorted by the first UMAP component. Color scale indicates class prediction same as in D. **F:** Bar charts showing proportion of dendritic type per majority classifier prediction.

We used the mouse classifier to predict the different cell classes (C1, C2, C3) in the marmoset. We assessed the classifiers’ performance using a test set of 48 marmoset cells not included in any of the previous analyses but with available morphological information which was used to provide “ground truth” labels, as follows: fast spiking neurons (FSI) with congruent (i.e., basket cell) morphology were determined to be C1 cells, whereas other aspiny neurons that could be identified as different morphological types (such as bi-tufted cells, NGFs, etc.) were determined to be C2 cells (see **Fig 2A,B**). C3 cells were drawn from the pool of all spiny cells. Marmoset cells were classified with a median accuracy of 83.3% (**Fig 3B**). Confusion matrices provided insight into classification accuracy by class (**Fig 3C**). Compared to mouse, erroneous classification of C1 and C3 as C2 cells was more common (positive predictive values: mouse = 78.7% vs marmoset = 55.7%). Marmoset C1 and C3 cells were identified with high specificity (positive predictive value: C1 = 94.3% and C3 = 95.8%).

We next examined the correspondence between the categories identified by the classifier and dendritic type, as an additional evaluation independent of intrinsic membrane properties (**Fig 3E-F**). Final class labels were determined by the most frequent prediction throughout different rounds of classification (repetitions) with different training subsamples (**Fig 3E**). Composition of dendritic type by putative class (**Fig 3F**) is in line with positive predictive values of the test data sets. In marmoset, 99% of putative C3 cells were spiny. The fraction of aspiny cells in the putative C2 cells (68.1%) matches very well with the share of ground truth interneurons (i.e., C1 and C2) in the totals of the test data (68.8%). In summary, these findings indicate that the cross-species classification leads to reliable labels for both C1 and C3, but less reliable labels for C2.

We repeated the same procedure in a sample of human neurons obtained from the AIBS database and the results were qualitatively similar for the different classes. However, there was a slight decrease in performance relative to the marmoset (**SFig 3D**).

### Comparison of intrinsic properties between V1 and LPFC neurons

We used the classifier to identify C1 and C3 cells and then compared the intrinsic properties of these classes between V1 and LPFC. First, we focused on C3, which includes EXC cells. A summary of comparisons in key features is depicted in **Table 1**. We focus on supragranular (layers II-III) neurons because these neurons process visual information from upstream areas (Roussy et al., 2022a, 2022b), and encode visual long term memories (Corrigan et al., 2022; Rouzitalab et al., 2023; Abbass et al., 2024), so they were intentionally targeted in our recordings (**Fig. 1D**).

We found that C3 cells from LPFC had a lower input resistance (p < 1e-4, effect size r_rb_ = 0.62), higher rheobase (p < 1e-4, r_rb_ = -0.63), and less rectification (p < 1e-4, r_rb_ = -0.64) than V1 cells (**Fig 4A, B, Tab1**). These results suggest that LPFC EXC cells are less excitable than V1 cells. To investigate the factors that may underlie the differences between areas, we examine how the size of C3 dendritic trees (quantified by total length) relates to the cell’s input resistance (**Fig 4C**). Indeed, both parameters showed a correlation of rho = -0.929 (p-value < 0.001), meaning that cells with shorter dendritic trees have a higher input resistance. V1 C3 cells had systematically smaller dendritic trees than LPFC C3 cells. V1 cells also fired significantly narrower action potentials with a robust effect size (p < 1e-4, r_rb_ = -0.42; **SFig 5**). Interestingly, firing rate adaptation, the increase in inter-spike interval duration after stimulation onset, was larger in LPFC than in V1 (p = 0.0032, r_rb_ = -3.14). A list of comparisons for several intrinsic features appears in **Tab1**. A total of 14 out of 19 features showed significant differences between areas.

**Figure 4:**
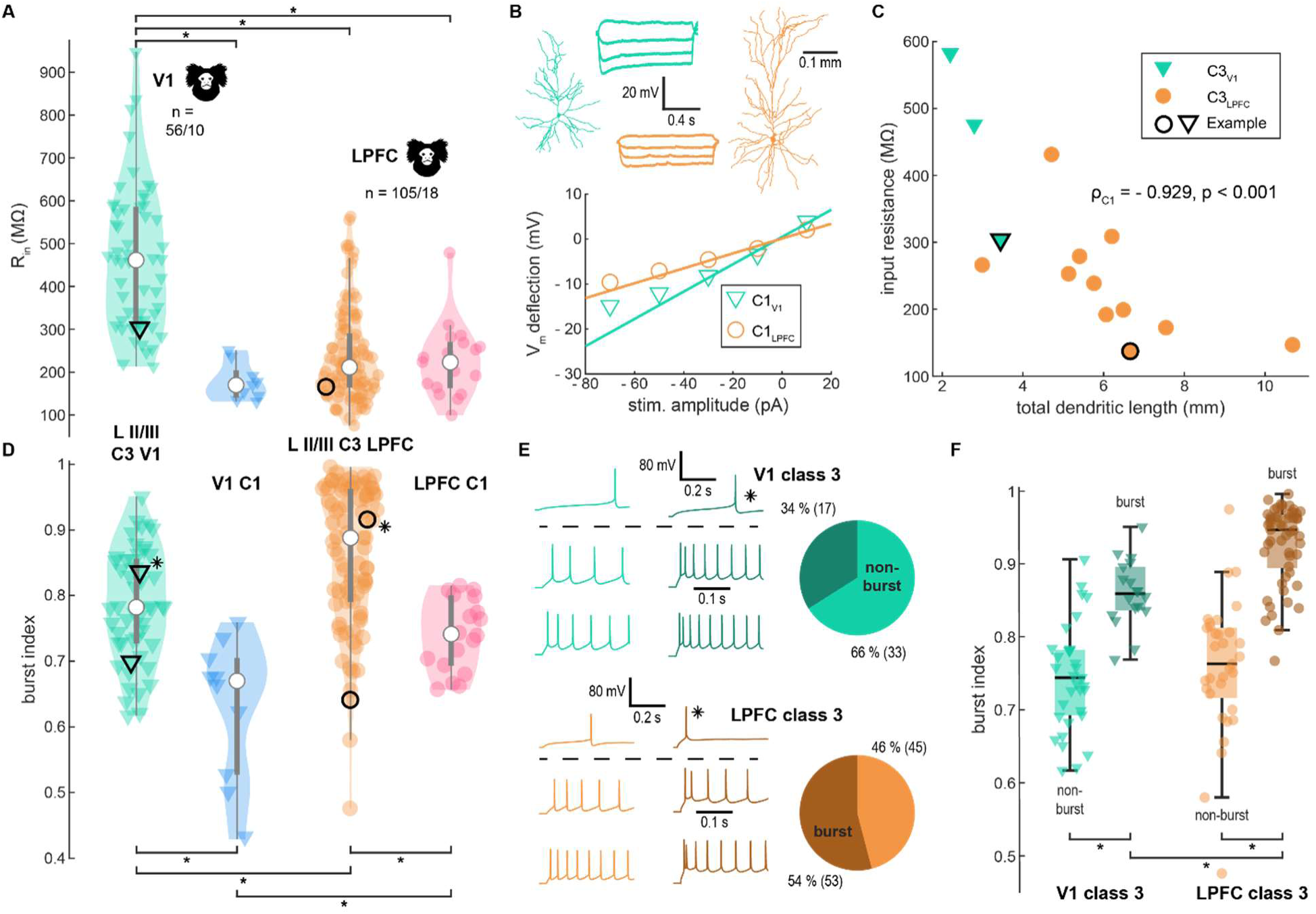
Input resistance and bursting across areas. **A, D:** Violin plots of class 3 (V1, turquois; PFC, orange) and class 1 (V1: blue, LPFC: pink) for input resistance (A) and burst index (D). Black contours signify example cells (triangle: V1, circle: LPFC) shown in B (for A) and E (for D). Asterisk in D labels bursting class 3 neuron. **B:** Above: Dendritic morphology of two example of class 3 cells (LPFC: orange, circle in A; V1: turquois, triangle in A) together with 4 subthreshold traces. Below: UI curve showing how input resistance of cells above was determined as the slope of linear fit of three lowest subthreshold stimulation. **C:** Total dendritic length and input resistance strongly correlate in class 3 cells. **E:** Example of bursting (V1, dark green; LPFC, brown) and non-bursting (V1, turquois; LPFC, orange) class 3 cells: Sweeps above the dashed line are from rheobase and show the first 0.5 s, instead of the first 0.25 s. Bursting is marked by doublets at medium to high stimulation, but not on rheobase level. **F:** Comparison of burst index of class 3 cells split by burst/non-burst distinction.

We further compared intrinsic properties of C1 fast spiking inhibitory interneurons between the two areas (see **STab 1**). Out of 19 analyzed features, 4 showed significant differences indicating that intrinsic features in interneurons were more similar between areas. We found that AP width was smaller in V1 than LPFC (p = 0.0369; r_rb_ = -0.51). The latency at the herosweep (see **methods** and **STab 2**) was longer in LPFC than in V1 (p = 0.0369, r_rb_ = -0.58). The difference in trough was also significant with a more negative value in V1 than in LPFC (p = 0.0369, r_rb_ = - 0.54). The rheobase showed a large difference and effect size between areas (see **STab1**), but it did not reach statistical significance.

### Burst firing in C1 and C3 cells

Bursts are short, high frequency trains of action potentials or spikes that could elicit a response in the postsynaptic cell with higher probability than isolated spikes (Zeldenrust et al. 2018). We computed burst indices in all the recorded neurons (see methods). Both C1 and C3 neurons showed an increased burst index in LPFC relative to V1 (**Fig 4D**; C1: p = 0.0369, r_rb_ = -0.51; C3: p < 1e-04, r_rb_ = -0.39). Some C3 neurons in both areas fired duplets of action potentials at the beginning of stimulation at medium to high intensities (**Fig 4E**). Violin plots show a distribution that was best fitted by the sum of 2 Gaussians rather than by a single Gaussian function (LPFC data, Akaike Information Criteria, AIC 1 Gaussian = -140; AIC 2 Gaussians = -181), indicating bimodality. Consequently, we divided all C3 cells into two subpopulations according to the presence of duplets, or two-spikes bursts (see example traces in **Fig. 4E**). After isolating C3 bursting cells we found that: 1) bursting cells were more frequent in LPFC than in V1 (54% vs 34%, **Fig 4E**), and 2) bursting cells in LPFC have a higher burst index than bursting cells in V1 (p < 1e-4, r_rb_ = -0.50, **Fig 4F**). In summary, C3 LPFC neurons show an increase in bursting relative to V1.

Interestingly, some of the C1 neurons in LPFC also showed a significant increase in burst index (**Fig 4D**), which was associated with a bursting fast-spiking phenotype (purple traces in **Fig 5A).** Unlike C3 neurons, C1 cells responded with bursts of multiple action potentials at rheobase stimulation, which was often accompanied by other features of intrinsic bursting at higher stimulation intensities such as sustained depolarization and varying AP wave forms. These bursting C1 neurons were exclusive to LPFC (5 out of 19, i.e., 26.3 %, **Fig 5B**) and appeared to be basket cells (BCs, see **Fig 5C**). With increasing stimulation, the gap in inter-spike intervals between the initial burst and post-burst narrowed, making it harder to distinguish bursting vs non-bursting fast spiking cells. At medium stimulus intensity, such as at the hero sweep, bursting C1 cells (purple color) had a higher burst index relative to other C1 LPFC cells (pink color, **Fig 5D**). LPFC cells we considered as not intrinsically bursting (pink color), show a higher burst index than their V1 counterparts (blue color).

**Figure 5:**
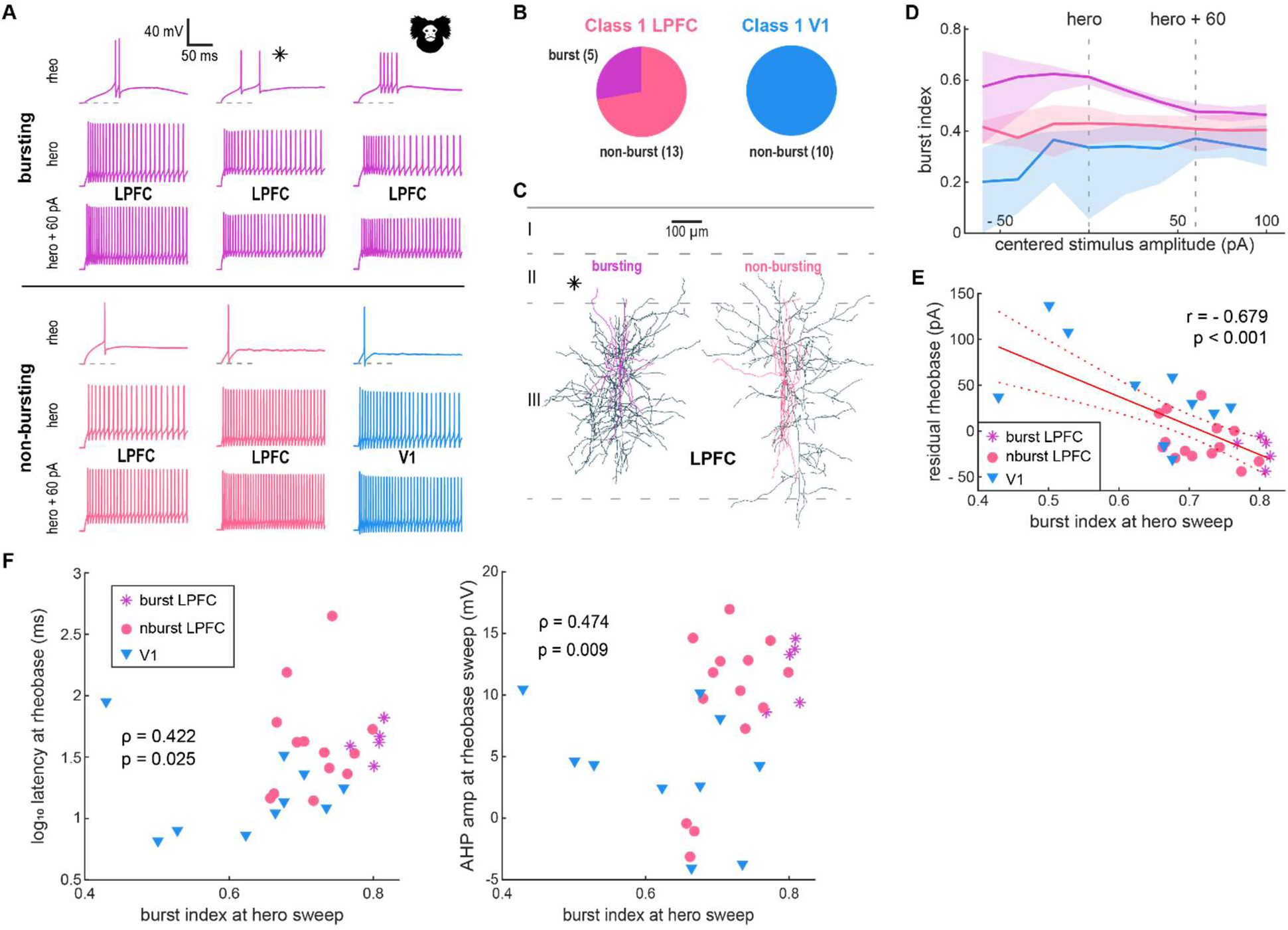
A bursting phenotype in C1 fast spiking basket cells. **A:** Example of various firing patterns of class 1 cells: neurons were divided into groups depending on bursting or lack thereof and cortical area: Bursting is marked by multiple spikes at rheobase level. **B:** Number of burst-type fast spiking basket cells among class 1 neurons in LPFC and V1. Bursting was observed in 28 % of LPFC class 1cells (purple vs pink) and not found in V1 (blue). **C:** Example morphology of bursting and non-bursting class 1 cells in LPFC. The dendritic tree is shown in purple (bursting) pink (non-bursting) while the axon is shown in dark blue. The asterisk corresponds to the example traces in A. Both groups are associated with basket cells. **D:** Burst index across stimulus intensity by subpopulation (burst-LPFC/non-burst-LPFC/V1). Line indicates median, shaded area indicates the IQR. **E:** Burst index negatively correlates with residual rheobase after regression with input resistance and difference between AP threshold and resting membrane potential. Markers are blue triangles for V1 cells, pink circles for non-bursting LPFC and purple stars for bursting LPFC cells. **F:** Burst index correlates with logarithm of latency and depolarized trough of the rheobase AP. **G:** Box charts comparing both logarithm of latency (left y-axis) and AP trough (right y-axis) by cortical area.

Intrinsic bursting cells typically have a prolonged active depolarization mechanism that facilitates the generation of action potentials. To explore this issue, we next examine the relationship between burst index and variables potentially linked to neuronal excitability. We first developed a linear regression model with rheobase as dependent variable and the ratio between the difference in the action potential threshold and the resting membrane potential (Dvoltage), and the input resistance (R) as predictor variable. This model (rheobase = B0 + B1*(Dvoltage/R) + residuals) captures the passive/ohmic aspect of the rheobase and provides a reasonable goodness of fit (adj. R2 = 0.73, p-value = 2.75e-09; model parameters [B0 = -3.654, p = 0.82597], [B1 = -826.86, p = 3.0106e-9*]). The model equation was used to obtain predicted values and residuals.

Negative residuals of this model reflect active depolarization mechanisms. We found a significant negative correlation between these residuals and burst index ( r = -0.642, p < 0.0001, see **Fig 5E**). These results suggest that an active depolarization mechanism underlies bursting in C1 cells. Burst index also correlated significantly with other variables, i.e., latency of the first action potential and amplitude of the afterhyperpolarization (latency p = 0.017, rho = 0.441; AHP amp , p = 0.009, rho = 0.474; see **Fig 5F**). Cells with higher burst index showed longer latencies to first spike at rheobase and weaker afterhyperpolarization, consistent with a depolarizing mechanism that activates slowly and persist beyond the first action potential within a burst.

### Differences in morphology between V1 and LPFC neurons

We investigated morphological features of C1 and C3 cells in LPFC and V1. We reconstructed 4 V1 and 11 LPFC pyramidal cells (PC). Two representative examples of PCs from V1 and LPFC are shown in **Fig 6A**. We focused on dendrites from PCs because their long axons are more likely to be truncated following slice preparation (Bruno et al., 2009; Oberlaender et al., 2011; Mohan et al., 2015; Goriounova et al., 2018). The total dendritic length of V1 PCs was half of their LPFC counterparts (median_LPFC_ = 5760 µm, median_V1_ = 3119 µm, p = 0.0352, r_rb_ = -0.63). Quantitative analysis showed that both apical dendrite length and complexity was significantly higher for PCs in LPFC than in V1 (apical length: p = 0.0352, r_rb_ = -0.63, apical complexity: p = 0.0352, r_rb_ = -0.63, **Fig 6C**). The result followed the same trend for basal dendrites with effect sizes of 0.42 and 0.52, although it did not reach statistical significance at 0.05.

**Figure 6:**
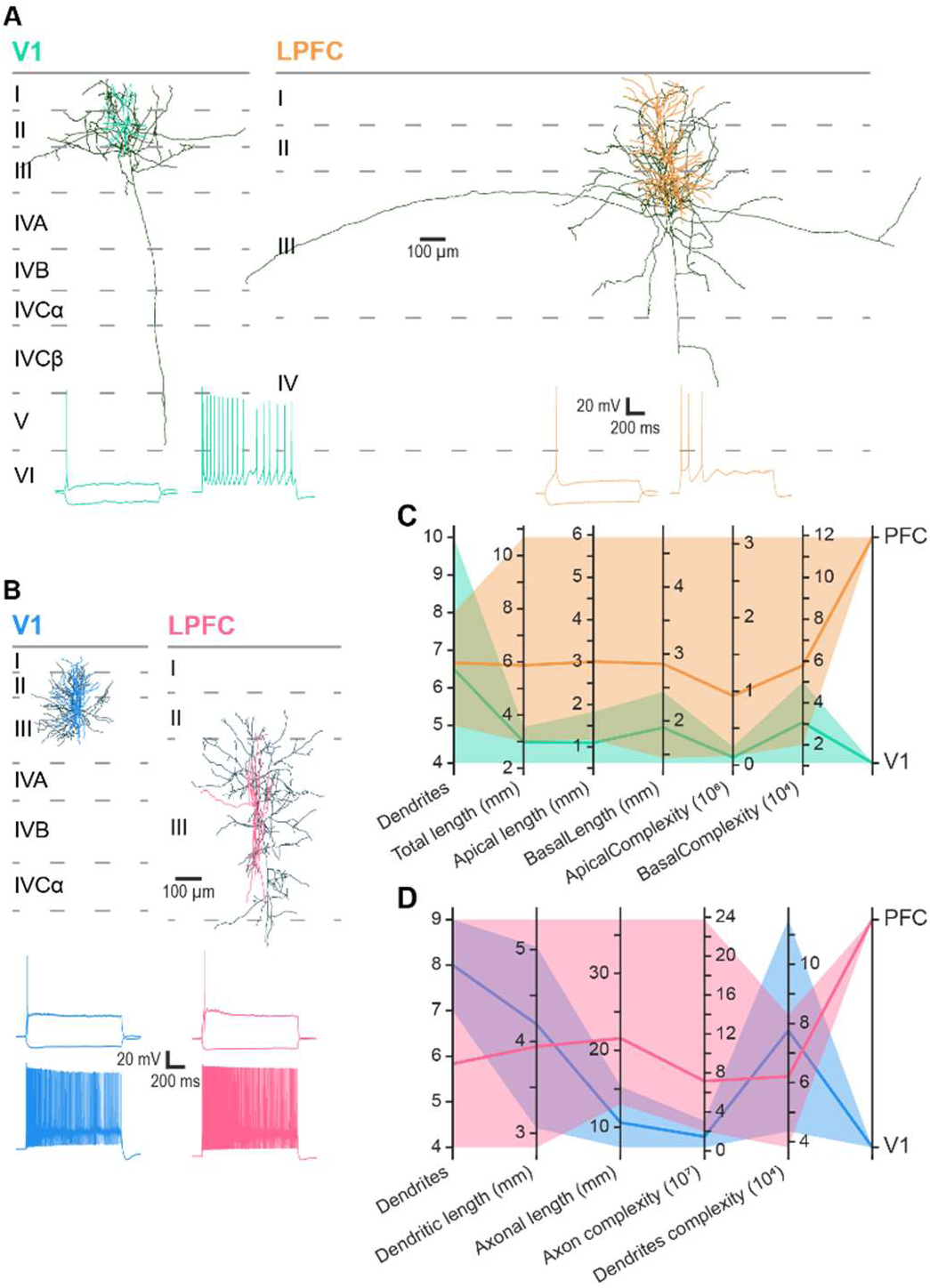
Comparison of morphology of pyramidal cells and fast spiking basket cells across areas. **A, B:** Examples of reconstructed marmoset neurons for area (left: V1, right: LPFC) and cell type of interest (A: pyramidal cells, B: fast spiking interneurons). Dendrites of pyramidal cells are colored in a lighter shade of orange for LPFC and in turquoise for V1, while the axons are in dark green. Dendrites of fast spiking basket cells were colored in pink for LPFC and light blue for V1, while axons are in dark blue. Below each neuron is the corresponding hero, rheobase and subthreshold response to the square-pulse stimulus. **C, D:** Parallel plots of key morphological quantifications for pyramidal neurons (C) and fast spiking basket cells (D). Transparent areas represent the range of the morphological feature for PFC and V1 respectively, while lines show the average for each brain region.

We also reconstructed 9 C1 (3 V1, 6 LPFC) fast-spiking basket cells (BC), all in layer III. Two example cells from V1 and LPFC are shown in **Fig. 6B**. In contrast to PCs, interneurons have a more localized axonal arborization, less likely to be affected by the thickness of the slice preparation. The axonal length of LPFC fast spiking BCs doubled the one of V1 neurons (median_LPFC_ = 19.743 mm, median_V1_ = 9.172 mm). There was over 5-fold increase in axon complexity in LPFC relative to V1 (median_LPFC_ = 52,0e6, median_V1_ = 9,2e6, **Fig 6D**). A larger sample of reconstructed cells would be needed to more systematically assess differences between areas.

In summary, we found that PCs in LPFC have larger dendritic complexity and length than in V1. On the other hand, a small sample of reconstructed fast-spiking BCs in LPFC show a more extensive and complex axon than in V1. These findings suggest that morphological changes have accompanied changes in intrinsic electrophysiological properties observed between V1 and LPFC neurons.

## Discussion

We compared electrophysiological and morphological features of single neurons in two different areas of the common marmoset visual cortical processing streams: V1, the input and earliest sensory area, and the LPFC, located far downstream at the end of the visual processing stream (Felleman and Essen, 1991). We conducted intracellular recordings using whole-cell patch-clamp in brain slices and anatomical reconstructions of neurons in both areas. To identify cell types, we built a random forest classifier using anatomical and transgenic labels, with electrophysiological features as predictors. We used the classifier to identify three classes of cells: C1 (aspiny fast-spiking interneurons), C2 (aspiny non-fast spiking interneurons), and C3 (spiny excitatory cells). We focused on C1 and C3 groups and compared neuronal intrinsic features between areas V1 and LPFC. C3 cells in V1 were more excitable, smaller, and fired narrower spikes than in LPFC. C1 neurons in the LPFC had longer latencies and a more depolarized action potential trough than V1 cells. In both C3 and C1 groups, intrinsic bursting was more commonly found in LPFC relative to V1 neurons.

### A tool for classifying neuron types using intrinsic properties and morphology

Neuronal diversity has been a puzzle for neuroscientists since the times of Santiago Ramon y Cajal, who described different cell morphologies across brain regions and species and suggested a relationship between the structure/morphology and function of a neuron (Szentágothai, 1990). Indeed, morphology and intracellular electrophysiological tools have been used to classify neurons (Ascoli et al., 2008). Progress in transcriptomics has advanced this field considerably (Saunders et al., 2018; Krienen et al., 2020; Yuste et al., 2020). However, studies of cell types have been mainly conducted in mice, where transgenic tools for single neuron classification are available and one can use large number of animals. In primates such as the macaque monkey some studies have examined electrophysiological and morphological features of pyramidal and non-pyramidal neurons (Gonzalez-Burgos et al., 2000; Zaitsev et al., 2005, 2012; Povysheva et al., 2007, 2008; Amatrudo et al., 2012; Luebke et al., 2015; Medalla and Luebke, 2015; Gilman et al., 2017; Medalla et al., 2017). However, in general they had low sample sizes and/or were limited to one cell type and/or brain area. More recent studies in resected tissue from human patients with epilepsy or brain tumors have also documented intrinsic features of neurons and their divergence from those of in the mouse brain (Kalmbach et al., 2018; Berg et al., 2021). However, they have also been restricted to one brain region, the temporal lobe, and changes due to the underlying pathology may have influenced their results.

Over the last decade, the common marmoset has emerged as a model in neuroscience research (Cyranoski, 2014; Mitchell et al., 2014; Mitchell and Leopold, 2015; Miller et al., 2016; Servick, 2018) opening a door to explore cell types in primate brains. In the current study we took advantage of the marmoset model to explore cell type diversity in areas V1 and LPFC of the primate cortical visual pathways. We used the same protocols as the AIBS to minimize technical and methodological confounds that could hinder comparisons of neuronal intrinsic properties and morphology between datasets (e.g., our marmoset data vs. AIBS mouse data). Our classification using transgenic mouse labels, as well as anatomical features (e.g., spiny and aspiny) resulted in three main classes of neurons (C1, C2, and C3) that include known cell types (Group et al., 2008; Tremblay et al., 2016; Zhou et al., 2020). C3 were mainly excitatory pyramidal cells, C1 fast spiking interneurons, and C2, more variable in their electrophysiological features, was mainly comprised of VIP and SST positive neurons in the mouse (Group et al., 2008; Defelipe et al., 2013; Tremblay et al., 2016; Merino et al., 2019; Zhou et al., 2020). This latter group likely included CB and CR positive interneurons in primates, thought to be homologues of SST and VIP positive cells in the mouse (Condé et al., 2004; Zaitsev et al., 2009, 2012; Torres-Gomez et al., 2020).

Our classification tool could be used by studies in humans and non-human primates that investigate the properties of different neocortical cell types when transgenic labels are not available. One important contribution of our study is highlighting homologies and differences between broad categories of cell types across species, which is critical to the use of animal models in research (**Fig. 2C, D)**. The differences between areas observed here agrees with the finding of transcriptomics studies in marmosets (Krienen et al., 2020). One general limitation of the existing maps is that the samples are biased to few brain areas (e.g., V1 in mice and Temporal cortex in humans (Gouwens et al., 2019), V1 and LPFC in marmosets (this study), which makes it difficult to perform wide inter-area comparisons across species. Future work is needed to fill this gap and provide a comprehensive map of functional and anatomical properties of neurons across different cortical areas of primates.

### Differences in intrinsic properties and morphology between cell types in V1 and LPFC

A main goal of this study was to identify distinctive features of cell types in areas V1 and the LPFC of the common marmoset. V1 and LPFC are in ‘opposite’ extremes (beginning and end) of the visual processing cortical hierarchy. In the initial stages of visual processing V1 neurons must fast and reliably process incoming information from the thalamus. We found that C3 excitatory cells in V1 showed high excitability and narrow action potentials, which may contribute to the elevated firing rates and short spike latencies relative to downstream areas reported by extracellular studies in macaques (Schmolesky et al., 1998; Reynolds et al., 1999). V1 also has the highest neuronal densities along the visual cortical pathways. The small C3 neuronal sizes may allow ‘packing’ more cells within the area. Indeed, excitatory V1 neurons receive highly spatially-specific LGN inputs corresponding to a small region of the retina, resulting in a high-resolution map of the visual field (Sincich and Horton, 2005). We also found that fast spiking interneurons (C1) in V1 show sustained high firing rates with low adaption in response to square current pulses. Additionally, V1 excitatory cells (C3) had high excitability, small dendritic trees and less intrinsic bursting. These features make V1 neurons suitable for linearly or quasi-linearly encoding important spatiotemporal details of visual inputs.

LPFC pyramidal neurons have large dendritic arborizations and many spines that can integrate inputs from a variety of sources (Young et al., 2014; Preuss and Wise, 2022). Pyramidal cells in the LPFC are known to fire with lower rates than in early sensory areas (Lennert and Martinez-Trujillo, 2013). Moreover, neurons output (i.e., firing rates) in LPFC has a higher correlation with performance than in upstream areas such as MT (Mendoza-Halliday et al., 2014) and presumably V1. Thus, LPFC signals seem to be more directly linked to our perceptions and actions than V1 signals. We found that excitatory cells (C3) in LPFC fired fewer spikes with the same current stimulation levels as V1. The lower excitability of LPFC pyramidal neurons is suited for selecting strong inputs corresponding to stimuli that are salient and/or behaviorally relevant and for producing sparse codes that may underlie our perceptual capabilities and sense of awareness (Panagiotaropoulos, 2024). However, the same structural and functional complexity may make LPFC pyramidal cells vulnerable to new categories of disease that would affect the highly developed cognitive skills of primates such as Schizophrenia (Glantz and Lewis, 2000; Lewis and González-Burgos, 2008; Arnsten, 2011) and Alzheimer disease (Arnsten et al., 2012).

Most interneurons receive their main inputs from local pyramidal cells. Pyramidal neurons and fast spiking interneurons intrinsic properties in V1 and LPFC may have diversified to perform computations according to the area’s input-output landscape. Indeed, certain types of interneurons are more abundant in LPFC than in early sensory areas (Condé et al., 2004; Torres-Gomez et al., 2020). One issue that remains unclear is when and how such diversification occurs. For the case of LPFC it is possible that the protracted development of primate species provides a prolonged window to adjust mRNA and protein expression (Bakken et al., 2016) based on local feedback (i.e., levels of firing rate) and therefore produce variations of what has been considered canonical cell types of the neocortex. In species with shorter protracted periods such as mice, neuronal types may appear more serially homologous (Harris and Shepherd, 2015; Gilman et al., 2017) perhaps due to a mode limited time window for epigenetics factors to transform the neuronal diversity landscape. This issue may be highly relevant for the study of mental disease (Lewis and Sweet, 2009) and deserves further investigation.

### Intrinsic bursting as a distinctive feature of LPFC neurons

A burst is a short, high frequency trains of spikes with a higher probability than single spikes to elicit a postsynaptic spike via temporal summation of excitatory postsynaptic potentials (Zeldenrust et al., 2018). Bursts could arise by intrinsic mechanisms or by network dynamics (Zeldenrust et al., 2018). Intrinsic bursters usually need a slow depolarization mechanism on top of which action potentials are generated. Intrinsic currents such as T-type Calcium have been isolated in bursters (Jahnsen and Llinás, 1984; Williams et al., 1997). The intrinsic bursting found in our LPFC cells may be linked to differential expression of T-type currents. In some LPFC fast-spiking C1 interneurons, depolarizing current injections elicited depolarizing events with slow kinetics that led to long latencies of action potentials evoked at low rheobase suggesting that at least a group of fast spiking interneurons in LPFC, likely basket cells, may express these channels (**Fig. 4A**).

Bursting has been associated to a variety of functions such as increase in the efficiency of synaptic transmission (Lisman, 1997; Csicsvari et al., 1998), and modulation of synaptic plasticity by increasing temporal summation of EPSPs and Long-Term Potentiation (LTP) in postsynaptic terminals (Thomas et al., 1998). The increase in bursting in LPFC relative to V1 may be linked to the ability of LPFC neurons to flexibly become tuned to novel objects (Freedman et al., 2002; Fitzgerald et al., 2012; Martinez-Trujillo, 2022). One interpretation of our results is that neurons in early areas such as V1 consistently process basic visual features (high stability - low plasticity), which is necessary for encoding the same sensory information (e.g., color, orientation) in a stable manner, regardless of whether it belongs to familiar or novel objects. On the other hand, neurons in the LPFC rapidly learn new associations of these basic features (low stability - high plasticity), which is necessary when encountering novel complex objects. Previous studies have reported a gradient in the ratio of NMDA- to AMPA-receptors (Yang et al., 2018; Froudist-Walsh et al., 2023) along the hierarchy of visual processing with LPFC having a high amount of NMDA glutamate receptors compared to V1. Combined with intrinsic bursting, NMDA gradients can lead to the enhanced ability of LPFC to build selectivity for new categories or rules (Freedman et al., 2001; Rouzitalab et al., 2023; Abbass et al., 2024). Our results can be interpreted within the framework of the feature integration theory of perception and attention (Treisman and Gelade, 1980; Treisman, 1982; Humphreys, 2015), in which essential features pre-attentively processed in early visual areas such as V1 and are latter assembled by the mechanisms of attention in the LPFC (Martinez-Trujillo, 2022).

### Serial homology and neuronal diversity in the primate visual system

It has been proposed that in the mammalian neocortex, cell types follow the principle of *serial homology*: cell types across brain areas are variations on a common theme organized in the same basic pattern. Differences between serially homologous structures are typically quantitative rather than qualitative (Harris and Shepherd, 2015). However, much of the data supporting this hypothesis have been collected in a single species, the mouse, and in sensory areas (e.g., barrel cortex or V1, Magrou et al., 2024). Mice do not show the stark differences in the morphology of pyramidal cells between V1 and PFC (Gilman et al., 2016) and perhaps the principle of serial homology applies to a larger degree than in primates. Previous studies observed a strong difference in structural complexity, from branching patterns (Gilman et al., 2016) to increases in the number of dendritic spines (Arnsten et al., 2012; Yang et al., 2013) between V1 and LPFC neurons. Moreover, we see qualitative differences in firing patterns such as the emergence of intrinsic bursting in LPFC. Thus, primate brains with expanded neocortices seem to have diversified single neuron structure and function to cope with the demands of their ecological niches achieving more brain computational power and behavioral complexity and making them some of the most adaptable species in the planet.

Inhibitory interneurons, have been classically considered less diverse across areas and species (Tremblay et al., 2016). However, most of what we know about interneuron diversity is from studies in mice. Interestingly, a recent study has shown substantial variation in the expression of different types of mRNAs across various species (mice, ferrets, marmosets and human, (Krienen et al., 2020)). In agreement with our results, several transcripts were differentially expressed in primate interneurons depending on the area (Krienen et al., 2020), supporting the notion of existing qualitative differences in interneurons subtypes across areas in primates. Thus, if there is serial homology across different areas of the primate visual system, it may be restricted to some cell types. Neuronal diversity across brain areas may be a hallmark of primate evolution and may arise during long-protracted periods of brain development (Otani et al., 2016). How this diversification of cell types has impacted normal brain function and vulnerability to new categories of brain diseases affecting humans is an open question.

## Conclusion

Our study revealed several differences in intrinsic properties of pyramidal cells and fast spiking inhibitory interneurons between areas V1 and the LPFC. An important contribution of our study is to show that area specialization in the primate visual system permeates the most basic level of signal processing, the single neuron. The data corresponding to the manuscript are available as a resource (www.primatedatabase.com).

## Acknowledgements

We would like to thank all veterinarians, veterinary technicians and experimenters taking care of the animals used in the study. At Western University: Kristy Gibbs, Rhonda Kersten, Cheryl Vander Tuin, Jarrod Dowell, Diego Buitrago Piza, Tsz Wai Bentley Lo. In Goettingen: Corinna Boike, Diana Kaltenborn, Verena Arndt, Diana Kues, Martina Bleyer, Daniel Aschoff.

In addition, we would like to thank Irma Meteluch, Patricia Sprysch, Pavel Truschow for excellent technical assistance around tissue processing and imaging as well as numerous student assistants for manual reconstruction of neuronal morphology, in particular Rakshit Sharma, Abdirahman Mohamed Salat, Katharina Bodes, Ima Mansori, Nicolas Zdun, Marvin Luca Schmidt, Sabrina Schild, Sophia Heidenreich, Leander Isaak Matthes, Serena Lam, Luca Auge, Josi Dittmann, Omar Aljameh, Amirhossein Haddadbakhodaei.

Furthermore, we would like to thank other members of the NeuroNex grant for giving their time and feedback on discussion of the project, namely John Meng, Xiao Xing Wang, Takeaki Miyamae, Benjamin Corrigan.

Last, but not least, we want to show our gratitude to Eric Kuebler, Kartik Pradeepan and William Assis, who have created the website featured in this publication together with coauthor Sam Mestern.

This work was primarily supported by the internation research grant NeuroNex funded by the Canadian Institutes of Health Research (CIHR reference: FL6GV84CKN57, MF, MSJS, SM, SV, SE, SJT, WI, JMT) , the Deutsche Forschungsgemeinschaft (DFG, Projektnummer 436260547, SP, JR, FP, ST, AN, JFS) and the National Science Foundation (NSF, grant number: 2015276, AFTA, DAL, GGB). JMT and WI were also supported by the Natural Sciences and Engineering Research Council of Canada (NSERC) and BrainsCAN grants.

## Author Contribution Statement

M Feyerabend: data curation (lead), formal analysis (lead), investigation (lead in Canada), visualization (equal), writing – original draft (equal), writing – review & editing (support)

S Pommer: data curation (support), formal analysis (support), investigation (lead in Germany), visualization (equal), writing – original draft (equal), writing – review & editing (support)

M Jimenez-Sosa: data curation (support), investigation (equal) J Rachel: investigation (equal)

J Sunstrum: investigation (equal)

F Preuss: investigation (equal)

S Mestern: investigation (equal)

R Hinkel: resources (lead)

M Mietsch: resources (support)

S Viyajraghavan: resources (support)

S Everling: resources (support), funding acquisition (support)

S Treue: resources (support), funding acquisition (lead),

AFT Arnsten: funding acquisition (lead), project administration (lead), conceptualization (support)

DA Lewis: funding acquisition (support), writing – review & editing (support) F Wolf, S McCarroll: funding acquisition (support)

L Muller: funding acquisition (support), project administration (support)

F Krienen, J Murray: funding acquisition (support), conceptualization (support)

SJ Tripathy, G Gonzalez-Burgos: funding acquisition (support), conceptualization (support), writing – review & editing (support)

W Inoue, JF Staiger, A Neef : funding acquisition (support), resources (support), supervision (equal), writing – review & editing (support)

J Martinez-Trujillo: funding acquisition (lead), writing – original draft (equal) , writing – review & editing (lead), supervision (equal)

## Notes

https://primatedatabase.com/

### Competing Interest Statement

The authors have declared no competing interest.

### Summary of Updates

A change in author's list, formatting issue in the references

