## Supplemental tables and figures for "Single neuron diversity supports area functional specialization along the visual cortical pathways"

### Supplementary Figures

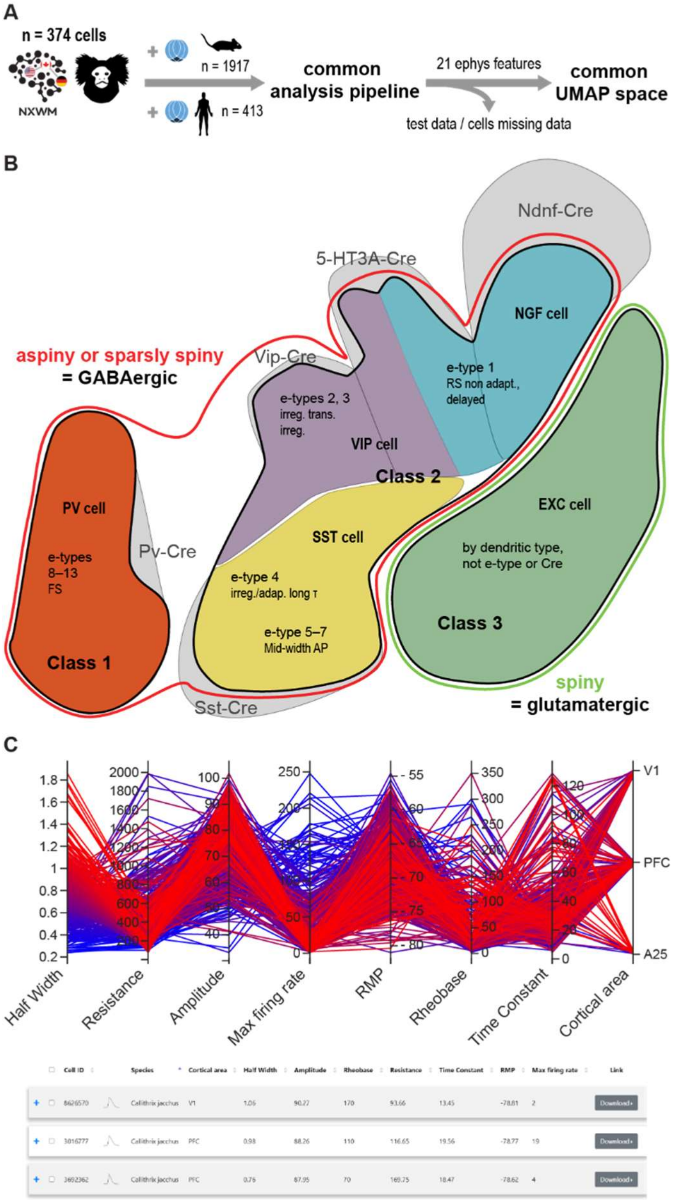


***Supplementary Figure 1: Processing pipeline and class selection criteria for the UMAP classifier.***
***A:*** *Processing pipeline of marmoset, mouse and human data. NEURONEX logo indicate the different member labs by country.* ***B:*** *Criteria for training data for class 1, 2 and 3 based on Cre-dependent mouse lines and e-types.*
***C:*** *Visualization of electrophysiological features across cells as seen on www.primatedatabase.com. Each line represents a cell. Color of the line was assigned by first feature, i.e. action potential width.*

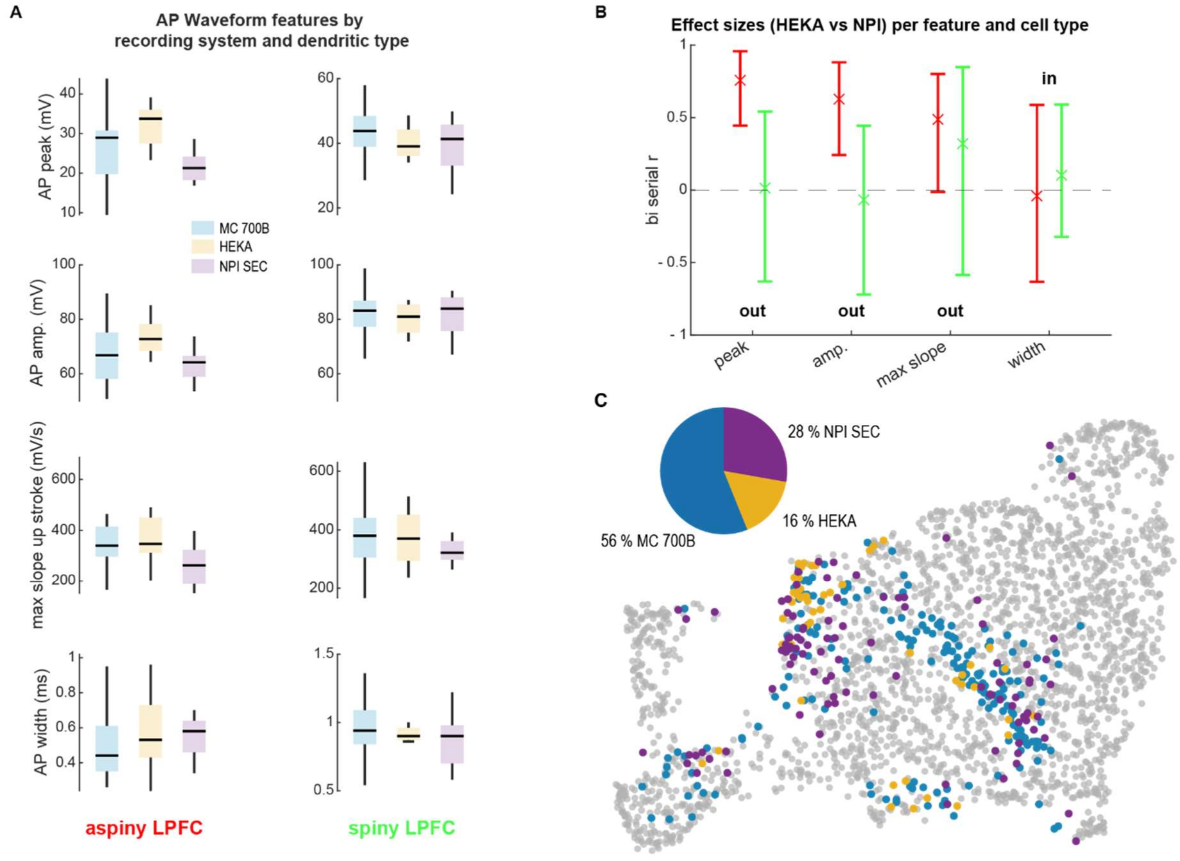


***Supplementary Figure 2: Effects of recording system on electrophysiological features in PFC.***
***A:*** *Comparison of AP waveform features by dendritic type (left: aspiny PFC, right: spiny PFC): in blue Multiclamp (MC) 700B (spiny: n = 56, aspiny: 15), in gold HEKA (spiny: n = 4, aspiny: n = 8 ), and in purple npi's single electrode clamp (SEC) amplifier (spiny: n = 12, aspiny: n = 9 ).* ***B:*** *Comparison of effect sizes across features with 95 % confidence interval between HEKA and NPI per feature and dendritic type (red: aspiny, green: spiny) based on the data shown in A. The first three features were excluded as input for UMAP projection due to a noticeable effect in aspiny cells caused by amplifier type.* ***C:*** *2D UMAP Projection with marmoset cells colored by recording system.*

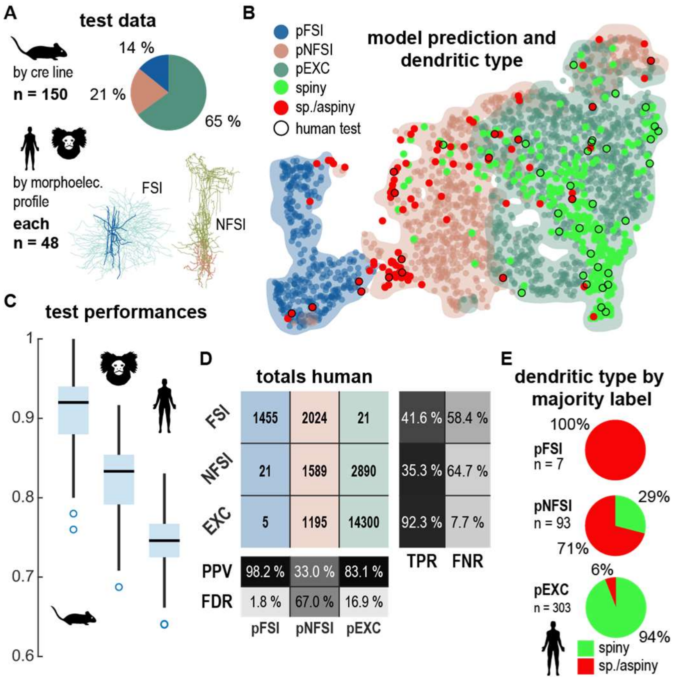


***Supplementary Figure 3: A:*** *Panel of pie charts showing cell type composition of training (mouse) and human test data. Three different classes were used: Class 1 (dark blue; C1), Class 2*
*(ocher, C2), with example morphologies of human C2 (right side, ocher dendrites) and human C1 cells (left side, dark blue dendrites). B: Example UMAP visualization of one model prediction referenced with dendritic type of human cells. Shaded areas indicate high density of the respective class. C: Performance of classifier across repetitions. Horizontal boxplots show distribution of performances. Median test performances by species were 91.3% for mouse, 83.3% for marmoset and 74.5% for humans. D: Confusion matrices for human test data showing all classification totals. The positive predictive value (PPV) for C1 cells is over 90% similar to the marmoset. FDR = False Discovery Rate, TPR = True Positive Rate, FNR = False Negative Rate. E: Pie charts showing proportion of dendritic type per majority classifier prediction.*

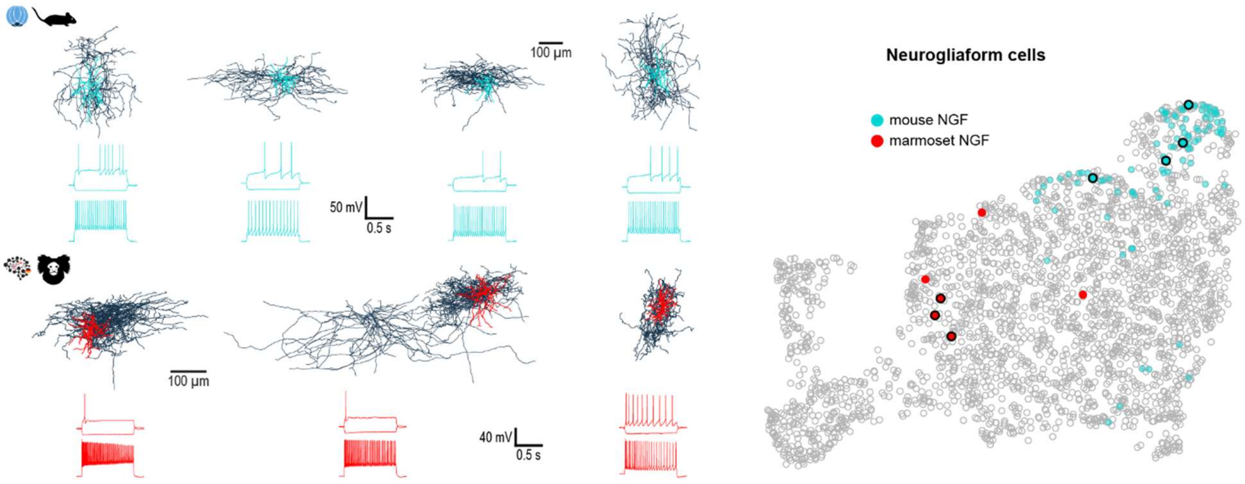


***Supplementary Figure 4****: Reconstructions of example NGFs from the mouse (AIBS, top) and marmoset (bottom). The dendrites are coloured in teal (mouse) and red (marmoset) with the axons in dark blue. Next to each reconstruction are the corresponding subthreshold, rheobase and hero sweeps. Note the early spike onset in the marmoset NGFs. The right side shows a 2D-UMAP space of mouse and marmoset cells with the localization of the NGF cells (mouse, teal circles; marmoset, red circles). The example reconstructions on the left side are marked with a black border. Scale bars equal 100 µm.*

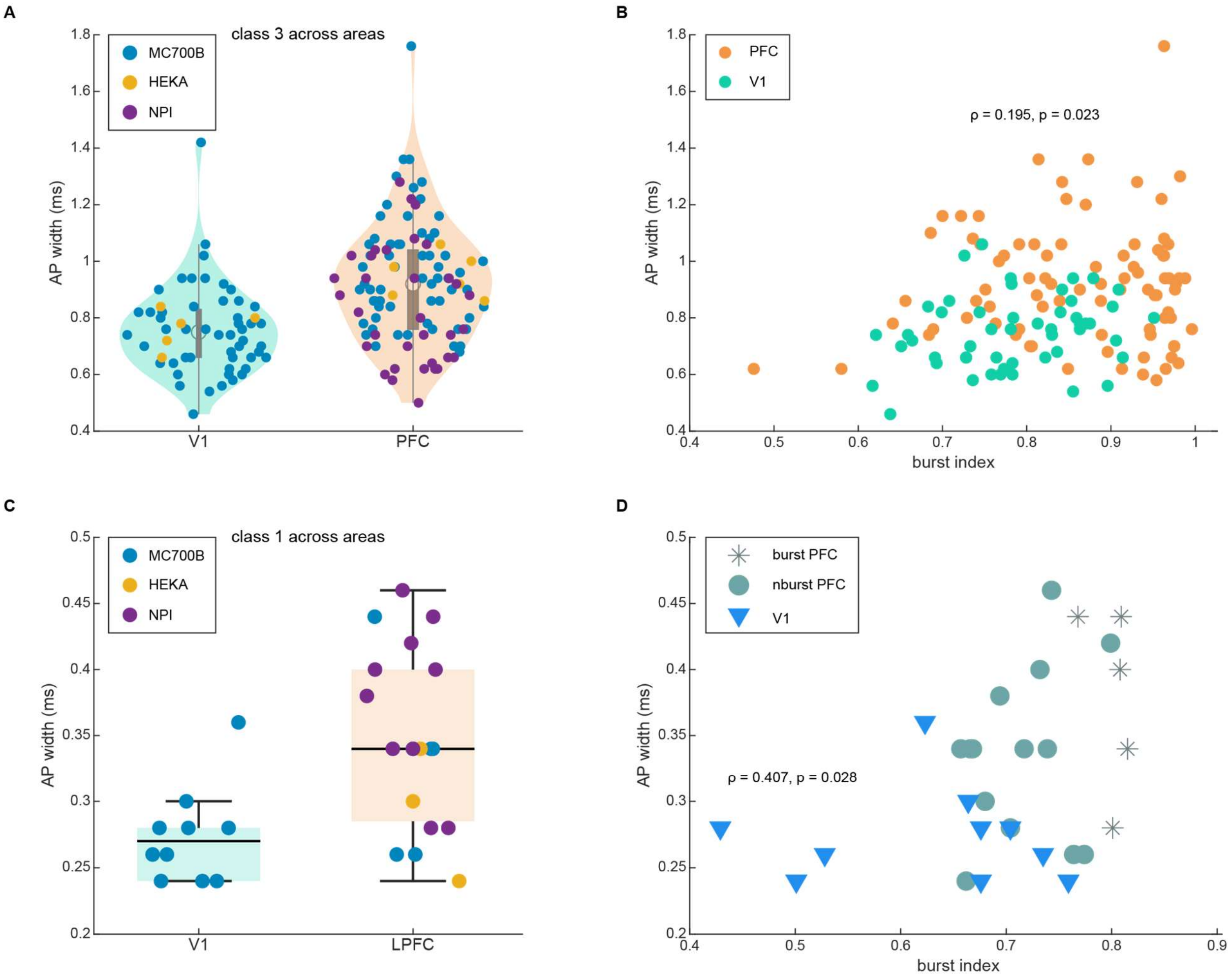


***Supplementary Figure 5: AP width difference between V1 and LPFC replicated across laboratories and relationship with bursting index.***

***A,C*** *Distribution of action potential (AP) width by area (turquoise : V1, orange: PFC): class 1 cells are depicted with box plots in C, whereas class 2 cells are depicted in violin plots. Marker color indicates different acquisition system (blue: MC700B, yellow: HEKA, purple: NPI).* ***B*** *Scatter plot of burst index and AP width for data shown in A. Features depicted correlate weakly (Spearman rho* $\rho = 0.195$*).* ***D*** *Scatter plot of burst index and AP width for data shown in C. Features depicted correlate moderately with* $\rho = 0.407$*.*

### Supplementary tables

***Supplementary Table 1: Electrophysiological characteristics of Class 1 neurons in LPFC and V1***

| Feature | LPFC (n = 10) | V1 (n = 19) | adj. p-value | z-value | Effect size |
|---|---|---|---|---|---|
| Vrest, mV | -66.33 | -63.93 | 0.2951 | -1.54 | 0.29 [-0.09;0.64] |
| $R_{inHD}$, MΩ | 231.32 | 222.46 | 0.3278 | 1.26 | -0.24 [-0.57; 0.13] |
| rheobase, pA | 85.00 | 195.00 | 0.2103 | -1.92 | 0.37 [-0.02; 0.70] |
| $\tau$, ms | 12.02 | 13.36 | 0.8080 | -0.30 | 0.06 [-0.32; 0.42] |
| sag ratio | 1.11 | 1.13 | 0.6959 | -0.60 | 0.12 [-0.29; 0.50] |
| inst. rectification | 0.82 | 0.81 | 1.0000 | 0.00 | 0.00 [-0.42; 0.43] |
| AP width | 0.35 | 0.27 | 0.0369* | 2.69 | -0.51 [-0.75; -0.22] |
| amp. fast trough | -21.03 | -24.48 | 0.3132 | 1.45 | -0.28 [-0.60; 0.06] |
| rate of rheo, Hz | 2.00 | 1.00 | 0.2942 | 1.61 | -0.31 [-0.59; 0.03] |
| med. inst. rate, Hz | 114.42 | 145.56 | 0.3278 | -1.31 | 0.25 [-0.15; 0.63] |
| IQR inst. rate, Hz | 48.87 | 47.43 | 0.6959 | 0.64 | -0.13 [-0.50; 0.28] |
| adaRat Blast/B1 | 0.55 | 0.44 | 0.3278 | 1.33 | -0.26 [-0.58; 0.12] |
| fIslope, Hz/pA | 0.66 | 0.78 | 0.8080 | -0.34 | 0.07 [-0.33; 0.43] |
| cvISI | 0.24 | 0.18 | 0.4412 | 1.03 | -0.20 [-0.56; 0.16] |
| rate of hero, Hz | 71.00 | 85.00 | 0.7869 | -0.44 | 0.09 [-0.35; 0.51] |
| latency at hero, ms | 7.78 | 4.58 | 0.0369* | 3.05 | -0.58 [-0.78; -0.31] |
| adaptation index | 0.01 | 0.00 | 0.2942 | 1.65 | -0.32 [-0.65; 0.08] |
| difference in trough | -1.92 | -4.81 | 0.0369* | 2.82 | -0.54 [-0.79; -0.21] |
| burst index | 0.74 | 0.63 | 0.0369* | 2.66 | -0.51 [-0.74; -0.21] |

Note: Values are median of electrophysiological characteristics in class 1 cells of LPFC and V1.

***Supplementary Table 2:*** *Description of electrophysiological features*

| Feature | Definition | UMAP input |
|---|---|---|
| AP width | Width of the 1st AP of the rheobase sweep at half amplitude determined from threshold to peak. | Yes |
| AP threshold | Threshold of the first rheobase action potential determined as membrane potential at which 5% percent of the peak slope of the rising phase is reached. | Yes |
| fast trough | Minimum membrane potential within 1.5 ms after the first rheobase AP has reached threshold level again. | Yes |
| slow through | Minimum membrane potential after the first rheobase AP has reached threshold level again up to the next AP or the stimulus end. | Yes |
| latency | Time difference between stimulus onset and threshold of the first action potential at the rheobase sweep. | Yes |
| rheobase rate | Number of spikes at the rheobase sweep. | Yes |
| hero sweep rate | Number of spikes at the hero sweep. The hero sweep is defined as the sweep closest to 65 % of the sweep with max. firing rate. | Yes |
| hero sweep current step | Current step of the hero sweep. | Yes |
| hero sweep latency | Time difference between stimulus onset and threshold of the first AP at the hero sweep. | Yes |
| median instantaneous rate | ISIs pooled across stimulus intensities. This is inverse to the median ISIs. | Yes |
| $P_{90}$ total ISIs | 90th percentile of ISIs pooled across stimulus intensities. | Yes |
| $P_{10}$ total ISIs | 10th percentile of ISIs pooled across stimulus intensities. | Yes |
| interquartile range total ISIs | Interquartile range of ISIs pooled across stimulus intensities. | Yes |
| adaptation ratio (last bin) | Stimulus divided into 13 bins of 77 ms each. Spike counts per bin are summed up across stimulus intensities. Ratio is calculated by first bin and last bin with non-zero value. | Yes |
| input resistance (highest deflection) | Slope of linear fit of IU data of the three lowest current steps. Voltage determined by membrane potential change to highest deflection within the first 200 ms of the stimulus. | Yes |
| input resistance (steady state) | As above but: Voltage determined by membrane potential change to steady-state potential within the last 200 ms of the stimulus. | Yes |
| time constant / $\tau$ | Maximum tau of hyperpolarizing current steps with a membrane deflection between 2 and 11 mV. Tau is calculated as time point when the exponential fit of the membrane potential deflection from stimulus onset to highest deflection reaches 66%. | Yes |
| $V_m$ | Mean of all sweep baseline membrane potentials of all sweeps calculated as mean of the prestimulus interval. | Yes |
| $V_m$ sag sweep | Mean of the prestimulus interval of the sag sweep. | Yes |
| delayed rectification | Ratio between steady-state membrane deflection and hypothetical steady-state deflection based on the input resistance at the most hyperpolarizing sweep. | Yes |

| | | |
|---|---|---|
| instantaneous rectification | Ratio between highest membrane deflection and hypothetical highest deflection based on input resistance at the most hyperpolarizing sweep. | Yes |
| rheobase | Current step at the rheobase sweep. Rheobase sweep is determined as the sweep with the lowest number of spikes. | Yes |
| sag | Difference in membrane potential between steady-state and highest deflection at the sag sweep. Sag sweep is calculated in the lowest hyperpolarizing stimulus sweep with a deflection higher than 11 mV. | Yes |
| sag ratio | (Sag + steady-state depolarization) divided by steady-state depolarization. Sag ration 1 = no sag. Sag ratio 1.5 = sag is half of the steady-state depolarization. | Yes |
| sag sweep current step | Current step at the sag sweep. | Yes |
| adaptation ratio ($2^{nd}$ bin) | As adaptation ratio (last bin) but: ratio is calculated by first and second bin. | Yes |
| trough difference | Difference in membrane potential between $1^{st}$ AP and $2^{nd}$ last AP in the hero sweep | Yes |
| trough ratio | Ratio between trough difference and difference between baseline membrane potential and trough of the $2^{nd}$ last AP. | Yes |
| peak adaptation | Ratio of the height of the $1^{st}$ AP and last AP of the hero sweep. | Yes |
| burst | 1-Ratio of $1^{st}$ ISI divided by mean ISI of the remaining ISIs from the hero sweep. | No |
| $CV_{ISI}$ | Coefficient of variation of ISIs of a sweep. Calculated as SD divided by mean at the hero sweep. | No |
| I-f slope | slope of a robust linear fit of the I-f curve. | No |
| adaptation index | Average rate of change in ISIs at hero sweep. | No |
